# Dopaminergic mechanisms underlying the expression of antipsychotic-induced dopamine supersensitivity in rats

**DOI:** 10.1101/2020.09.08.287664

**Authors:** Alice Servonnet, Florence Allain, Alice Gravel-Chouinard, Giovanni Hernandez, Casey Bourdeau Caporuscio, Mathilde Legrix, Daniel Lévesque, Pierre-Paul Rompré, Anne-Noël Samaha

## Abstract

Antipsychotic treatment can produce a dopamine-supersensitive state, potentiating the response to dopamine receptor stimulation. In both schizophrenia patients and rats, this is linked to tolerance to ongoing antipsychotic treatment. In rodents, dopamine supersensitivity is often confirmed by an exaggerated psychomotor response to d-amphetamine after discontinuation of antipsychotic exposure. Here we examined in rats the dopaminergic mechanisms mediating this enhanced behavioural response, as this could uncover pathophysiological processes underlying the expression of antipsychotic-evoked dopamine supersensitivity. Rats received 0.5 mg/kg/day haloperidol via osmotic minipump for 2 weeks, before treatment was discontinued. After cessation of antipsychotic treatment, rats showed a supersensitive psychomotor response to the D2 agonist quinpirole, but not to the D1 partial agonist SKF38393 or the dopamine reuptake blocker GBR12783. Furthermore, acute D1 receptor blockade (using SCH39166) decreased the exaggerated psychomotor response to d-amphetamine in haloperidol-pretreated rats, whereas acute D2 receptor blockade (using sulpiride) enhanced it. Thus, after discontinuation of antipsychotic treatment, D1- and D2-mediated transmission differentially modulate the expression of a supersensitive response to d-amphetamine. This supersensitive behavioural response was accompanied by enhanced GSK3β activity and suppressed ERK1/2 activity in the nucleus accumbens (but not caudate-putamen), suggesting increased mesolimbic D2 transmission. Finally, after discontinuing haloperidol treatment, neither increasing ventral midbrain dopamine impulse flow nor infusing d-amphetamine into the cerebral ventricles triggered the expression of already established antipsychotic-evoked dopamine supersensitivity, suggesting that peripheral effects are required. Thus, while dopamine receptor-mediated signalling regulates the expression of antipsychotic-evoked dopamine supersensitivity, a simple increase in central dopamine neurotransmission is insufficient to trigger this supersensitivity.

**HIGHLIGHTS:**

- Antipsychotic exposure can lead to a state of dopamine (DA) supersensitivity
- In rats, this DA supersensitivity potentiates d-amphetamine-induced locomotion
- We report that D2 transmission promotes DA supersensitivity and D1 transmission tempers it
- D-amphetamine’s central effects are also insufficient to reveal DA supersensitivity

## 1. Introduction

Antipsychotic drugs attenuate schizophrenia symptoms by blunting dopamine D2 receptor activity. However, long-term antipsychotic treatment can produce neuroadaptations that lead to supersensitivity to dopamine receptor stimulation. Antipsychotic-induced dopamine supersensitivity is linked to antipsychotic treatment failure and to an exacerbation of psychosis symptoms (Asper et al., 1973; MØller Nielsen et al., 1974; Chouinard et al., 1978; Margolese et al., 2002; Chouinard and Chouinard, 2008; Fallon and Dursun, 2011; Iyo et al., 2013; Gill et al., 2014; Chouinard et al., 2017). In rodents, a widely used behavioural index of antipsychotic-induced dopamine supersensitivity is a potentiated psychomotor response to d-amphetamine seen after cessation of chronic antipsychotic exposure (Smith and Davis, 1975; Rebec et al., 1982; Ericson et al., 1996; Meng et al., 1998; Samaha et al., 2007; Samaha et al., 2008; Carvalho et al., 2009; Bedard et al., 2011; El Hage et al., 2015; Servonnet et al., 2017). In this context, d-amphetamine serves as a pharmacological tool to probe the functional consequences of an acute increase in striatal dopamine release, as seen during psychosis (Howes et al., 2012). However, the anatomical location and nature of the dopaminergic effects through which d-amphetamine triggers a supersensitive behavioral response in antipsychotic treated rats are largely unknown. We investigated these effects here in rats with a history of chronic haloperidol treatment, as the answers could reveal underlying biological mechanisms and eventual therapeutic targets to suppress antipsychotic-evoked dopamine supersensitivity.

An important question concerns the role of dopamine-mediated neurotransmission. D-amphetamine stimulates dopamine, but also noradrenaline and serotonin transmission (Millan et al., 2002; Rothman and Baumann, 2003). Thus, we determined whether selective dopamine reuptake inhibition is sufficient to evoke a supersensitive response in antipsychotic-treated rats. Dopamine signals through dopamine D1-type and D2-type receptors. Selective D2 receptor stimulation evokes a supersensitive psychomotor response in antipsychotic-treated rats (Obuchowicz, 1999; Hashimoto et al., 2018), but whether D1 stimulation does the same is unknown. We addressed this here. As a complement, we also determined whether D1 and/or D2 receptor activity is required for the full expression of dopamine supersensitivity. At the neurobiological level, we assessed the effects of d-amphetamine on the activity of AKT/GSK3β- and cAMP/PKA-dependent signalling pathways, which are regulated by D1 and D2 receptors (Valjent et al., 2000; Beaulieu et al., 2004; Valjent et al., 2005; Beaulieu et al., 2007; Oda et al., 2015). This latter experiment suggested that dopamine supersensitivity is linked to increased dopamine transmission. Thus, we also examined the hypothesis that increasing ventral tegmental area (VTA) dopamine impulse flow is sufficient to trigger a supersensitive psychomotor response in haloperidol-treated rats. Lastly, we examined the contributions of d-amphetamine’s central effects, by determining whether injecting the drug into the cerebral ventricles is sufficient to trigger a supersensitive psychomotor response in antipsychotic-treated rats.

## 2. Methods

### 2.1. Animals

Male Sprague-Dawley rats (200-275 g; Charles River Laboratories, Montreal, QC) were used. In Experiments 1-4, rats were housed 2/cage. In Experiments 5-6, rats were housed 1/cage to avoid damage to intracerebral cannulae by conspecific. All rats were housed on a reverse dark-light cycle (lights off at 8:30 am). All testing took place during the dark phase. Water/food were available *ad libitum*. Experimental procedures were approved by the Université de Montréal’s ethics committee and followed the guidelines of the Canadian Council on Animal Care.

### 2.2. Drugs

D-amphetamine sulfate (Sigma-Aldrich, Dorset, UK), cocaine hydrochloride (Medisca Pharmaceutique, St-Laurent, QC), neurotensin acetate salt, DAMGO acetate salt (Bachem, Torrance, CA), (-)-quinpirole hydrochloride, SKF38393 hydrobromide and SCH39166 hydrobromide (R&D Systems, Minneapolis, MN) were dissolved in 0.9 % saline. Haloperidol (Sandoz, Boucherville, QC) was diluted in sterile water containing 0.5 % glacial acetic acid and pH was increased to ∼5 using NaOH. (-)-Sulpiride (Sigma-Aldrich, Milwaukee, WI) was dissolved in 0.9 % saline containing ∼1.4 % glacial acid acetic and pH was increased to ∼6.5 using NaOH. GBR12783 dihydrochloride (R&D Systems) was dissolved in DMSO, diluted in 0.9 % saline (final concentration of DMSO is 10 %) and pH was increased to ∼4 using NaOH. GBR12783 solubilised at pH above ∼4.5. Rats showed no visual/auditory signs of discomfort when receiving GBR12783 or its pH-matched vehicle. Apomorphine hydrochloride (Sigma, Oakville, ON) was dissolved in 0.9 % saline containing 0.1 % sodium L-ascorbate (Sigma, Oakville, ON). DAMGO, neurotensin and sulpiride solutions were frozen in aliquots and then thawed on testing days. GBR12783, apomorphine and its vehicle were prepared fresh on testing days. Systemic injections were given s.c., except cocaine and its vehicle, which were administered via intraperitoneal injections. Systemic injections were given in a volume of 1 mL/kg, except for GBR12783 and its vehicle (4 mL/kg) and SKF38393 and its vehicle (3 mL/kg). Microinfusions into the lateral ventricles or into the VTA were given in a volume of 2 or 0.5 µL/hemisphere, respectively.

### 2.3. Antipsychotic treatment

Standard antipsychotic treatment regimens used in the clinic achieve steady levels of striatal D2 receptor occupancy, when patients adhere to treatment (Farde et al., 1989; Remington et al., 2006; Mamo et al., 2008). To model this here, rats received haloperidol via an osmotic minipump (Alzet model 2ML2; Durect Corporation, Cupertino, CA) which produces continuous levels of D2 receptor occupancy during treatment (Kapur et al., 2003; Samaha et al., 2007), and control rats received sham surgeries (note that all behavioural tests took place after minipump removal, and so no rat had a minipump during testing). Rats received 0.5 mg/kg/day haloperidol. This dose achieves 73 % ± 14 SD striatal D2 receptor occupancy [unpublished observations, see (Kapur et al., 2003; Samaha et al., 2007)], and this is within the occupancy range that is therapeutically-efficacious in patients (Farde et al., 1992; Kapur et al., 1999; Kapur et al., 2000). Under isoflurane anaesthesia, minipumps were implanted subcutaneously (s.c.) for haloperidol-treated rats, and controls were sham-operated (Samaha et al., 2007). Seventeen days later, minipumps were removed, and controls were sham-operated again.

### 2.4. Intra-cerebral procedures

#### 2.4.1. Cannulae implantation

In Experiments 5-6, intra-cerebral cannulae were implanted at the same time as minipump implantation or sham surgery. Rats weighing 325-350 g were anesthetized with isoflurane (5 % for induction, 2-3 % to maintain anaesthesia) and placed on a stereotaxic apparatus. Rats received penicillin (3,000 IU, i.m.) and carprofen (1.5 mg, s.c.) at the beginning of surgery. A guide cannula (Experiment 5: 26 GA, model C315G; Experiment 6: 22 GA, model C313G; HRS Scientific, Montreal, Qc) was implanted in each cerebral hemisphere 2 mm above the VTA (A/P -5.9, M/L ±1.7, D/V -6.7, all mm relative to Bregma, M/L angle of 8°) or the lateral ventricles (A/P -1.1, M/L ±2.5, D/V - 3.3, all mm relative to Bregma, M/L angle of 10°). Four stainless steel screws were anchored to the skull and dental cement secured the cannulae. Guide cannulae were sealed with obturators (Experiment 5: model C315CD; Experiment 6: model C313CD; HRS Scientific).

#### 2.4.2. Intra-cerebral infusion

Microinfusions (0.5 µL/minute) were given via injectors protruding 2 mm beyond guide cannulae (Experiment 5: 33 GA, model C315I; Experiment 6: 28 GA, model C313I; HRS Scientific). The injectors were connected via tubing to 5-µL syringes placed on a microsyringe pump (HARVARD PHD, 2000: HARVARD Apparatus, Saint-Laurent, Canada). Following infusion, injectors were kept in place for an additional minute. On day 2 following minipump removal (before any behavioural testing), rats were brought to the testing room and were given an intra-cerebral infusion of 0.9 % saline for habituation. No behaviour was recorded.

#### 2.4.3. Histology

In Experiment 6, rats received an intracerebroventricular infusion of ink prior to brain extraction to facilitate histological verification. In Experiments 5-6, brains were frozen in isopentane and stored at -20 °C until processing. Placement of injector tips was determined on 40-µm coronal slices using the atlas of Paxinos and Watson (Paxinos and Watson, 1986). Data from rats with infusion sites outside of the targeted area were excluded from analysis.

### 2.5. Measurement of psychomotor activity

Psychomotor activity was measured using photocell counts and psychomotor activity ratings. Photocell counts—a measure of horizontal activity—were recorded in Plexiglas boxes (27 × 48 × 20 cm) equipped with 6 rows of photocells (3 cm above the box floor). An experimenter blind to condition rated psychomotor activity on minutes 5, 10, 20, 40 and 60 in Experiment 5 or every 10 minutes in Experiments 1-3 and 6 (unless an injection was given on minutes 30 or 60) using the following scale (Mattson et al., 2007) [modified from (Ellinwood and Balster, 1974)]: 1: asleep, 2: inactive, 3: normal in-place activity, 4: alert, rearing, normal level of locomotion, 5: rearing, high level of locomotion, 6: no rearing, normal/baseline level of locomotion, slow patterned behaviours (*e*.*g*., slow-patterned licking of cage walls), 7: no rearing, high level of locomotion, faster patterned behaviours (*e*.*g*., fast-patterned head rolling), 8: highly repetitive patterned behaviours in a restricted area and 9: backing up, abnormally maintained posture. A psychomotor activity rating ≥ 6 indicates stereotypy.

### 2.6. Western blot

Rats were briefly exposed to 5 % isoflurane and brains were extracted. Two-mm coronal slices were cut, and bilateral tissue punches were taken from the slice at ∼ +1.7 mm relative to Bregma in the nucleus accumbens, dorsal caudate-putamen, ventrolateral caudate-putamen and centromedial caudate-putamen using a 15-gauge sample corer. Striatal tissues were stored at -80°C until processing.

Striatal samples were mechanically homogenized in a lysis buffer (150 mM sodium chloride, 1 % triton X-100, 0.5 % sodium deoxycholate, 0.1% sodium dodecyl sulfate, 50 mM tris, pH = 7.4) containing protease and phosphatase inhibitors (Sigma-Aldrich, Oakville, ON). Homogenates were solubilized for 15 minutes on ice and then centrifuged at 16,000 g for 30 minutes at 4°C. The protein content of supernatants was measured using a BiCinchoninic acid Assay (BCA) protein assay kit (Thremo Fisher Scientific, Mississauga, ON, Canada). Equal amounts of protein in lysis buffer (10 µg) were dissolved in 25 μL of double distilled water (boiled at 95 °C for 5 minutes) containing loading buffer (4X; Bio-Rad Laboratories, Mississauga, ON, Canada) and reducing agent (20X; Bio-Rad Laboratories). Protein samples were loaded into a Bis-Tris 10 % pre-casted gels (Bio-Rad Laboratories). Proteins migrated for 60 minutes at 200 V and were then transferred to a polyvinylidene fluoride membrane (Bio-Rad Laboratories) for 2 hours (70 V, 4°C). Membranes were blocked in a solution of 5 % bovine serum albumin diluted in 0.1 % Tween 20/tris-buffered saline for 1 hour. Membranes were incubated overnight at 4 °C with the appropriate antibody: rabbit monoclonal anti-GSK3β (1:1,000; product # 9315, Cell Signalling Technology, New England BioLabs, Whitby, Ontario, Canada), rabbit polyclonal anti-p[Ser9]GSK3β (1:500; product # 9336, Cell Signalling Technology), rabbit polyclonal anti-AKT (1:1,000; product # 9272, Cell Signalling Technology), rabbit polyclonal anti-p[Ser473]AKT (1:500; product # 9271, Cell Signalling Technology), rabbit polyclonal anti-DARPP-32 (1:1,000; product # 2302, Cell Signalling Technology), rabbit monoclonal anti-p[Thr34]DARPP-32 (1:500; product # 12438, Cell Signalling Technology), rabbit polyclonal anti-ERK1/2 (1:50,000; product # 9102, Cell Signalling Technology), rabbit polyclonal anti-p[Thr202]ERK44/p[Thr204]ERK42 (1:10,000; product # 9101, Cell Signalling Technology) or mouse monoclonal anti-α-tubulin (1:50,000; product # T5168, Sigma-Aldrich, Oakville, Ontario, Canada). Membranes were then rinsed 4 times for 5 minutes with 0.1 % Tween 20/tris-buffered saline at room temperature. Membranes were incubated with the appropriate secondary antibody conjugated with horseradish peroxidase (goat anti-rabbit, 1:5,000 for phosphorylated kinase or 1:10,000 for total protein, product # 7074, Cell Signalling Technology; horse anti-mouse, 1:150,000, product # 7076, Cell Signalling Technology) for 1 hour at room temperature. Immunoreactive bands were revealed using the enhanced chemiluminescence reaction (Bio-Rad Laboratories) and the bands were placed against sensitive film (MidSci Scientific, Valley Parl, MI, USA) for a few seconds.

Densitometric levels were determined using Image Lab Software (Bio-Rad Laboratories). Background was subtracted for each band. The densitometric level of each band was normalized relative to the sum of densitometric levels across all tissue samples. Protein levels were then normalized relative to the corresponding level of the housekeeping protein α-tubuline. Protein levels were then normalized relative to the mean protein level of the control group that received saline injection prior to brain extraction. Using these values, we computed the ratio of phosphorylated protein levels over total protein levels.

### 2.7. Experiments

Fig. 1 illustrates experimental timelines. Locomotion tests started at least 3 days after haloperidol discontinuation and were given every 48 hours, 1 test/day, in counterbalanced order. Doses, routes of administration, treatment conditions and number of rats per condition (per control and haloperidol-treated group) for all experiments are detailed below and also summarised in Table I. Each experiment was undertaken in independent cohort of rats. We confirmed antipsychotic-induced dopamine supersensitivity in each cohort by measuring the psychomotor response to s.c. d-amphetamine (0 or 1.5 mg/kg). In Experiments 1-3, the different vehicles used produced similar locomotor responses and were pooled to form one vehicle control group.

**Table I.** Experimental parameters.

| Exp. | Agent(s) or vehicle and route | Dose |  | n per group | Design |
| --- | --- | --- | --- | --- | --- |
| **1** | S.c. or i.p. vehicle<sup>a</sup> | - | - | 29-31 | Mixed within/between-subjects, each rat received no more than 3 agonist doses (*i.e.*, 3 out of 6 possible doses) and 2 vehicle injections |
|  | S.c. GBR12783 | 5, 10 | mg/kg | 10-14 |  |
|  | I.p. cocaine | 2.5, 10 |  | 7-10 |  |
|  | S.c. apomorphine | 0.25, 0.5 |  | 9-10 |  |
| **2** | S.c. vehicle<sup>a</sup> | - | - | 16 | Mixed within/between-subjects, each rat received one dose of each agonist (*i.e.*, 2 out of 4 possible doses) and one vehicle injection |
|  | S.c. SKF38393 | 1, 10 | mg/kg | 8 |  |
|  | S.c. quinpirole | 0.15, 0.5 |  | 8 |  |
| **3** | S.c. vehicle & vehicle<sup>a</sup> | - | - | 31-32 | Mixed within/between-subjects, each rat received 4 combinations:<br>1) vehicle/vehicle<br>2) antagonist/vehicle<br>3) vehicle/d-amphetamine<br>4) antagonist/d-amphetamine |
|  | S.c. sulpiride & vehicle<sup>a</sup> | 25, 80 (sul) & vehicle | mg/kg | 7-8 |  |
|  | S.c. SCH39166 & vehicle<sup>a</sup> | 0.03, 0.1 (SCH) & vehicle |  | 8 |  |
|  | S.c. vehicle<sup>a</sup> & d-amphetamine | Vehicle & 1.5 (d-amph) |  | 31-32 |  |
|  | S.c. sulpiride & d-amphetamine | 25, 80 (sul) & 1.5 (d-amph) |  | 8 |  |
|  | S.c. SCH39166 & d-amphetamine | 0.03, 0.1 (SCH) & 1.5 (d-amph) |  | 7-8 |  |
| **5** | Intra-VTA vehicle | - | - | 7 | Within-subjects, all rats received neurotensin, DAMGO and vehicle injections |
|  | Intra-VTA neurotensin | 1 | nmol/hemisphere | 7 |  |
|  | Intra-VTA DAMGO | 0.3 |  | 7 |  |
| **6** | I.c.v. vehicle | - | - | 16-21 | Mixed within/between-subjects, one d-amphetamine dose per rat, all rats received vehicle |
|  | I.c.v. d-amphetamine | 50, 100 | µg/hemisphere | 6-10 |  |
| **1-6** | S.c. saline | - | - | 5-32 | Within-subjects, each rat received saline and then d-amphetamine, 30 or 60 min later |
|  | S.c. d-amphetamine | 1.5 | mg/kg |  |  |

<sup>a</sup>Rats receiving different vehicle solutions showed similar locomotor responses and were therefore pooled into one control group.

**Fig. 1.**
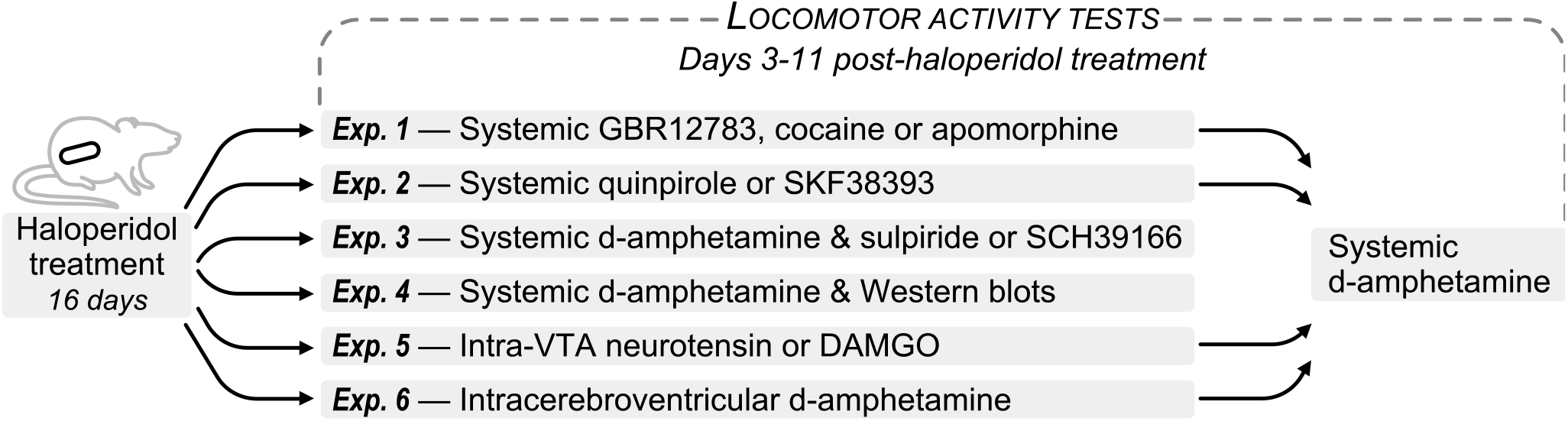
Experimental timeline. In all experiments, rats were given continuous haloperidol treatment (via a subcutaneously-implanted minipump set to administer 0.5 mg/kg haloperidol/day) for 16 days and testing took place 3 to 11 days following treatment cessation.

#### 2.7.1. Experiment 1: Stimulation of dopamine neurotransmission

We assessed whether selectively blocking dopamine reuptake with GBR12783 (Bonnet and Costentin, 1986) [0, 5 or 10 mg/kg (Le Pen et al., 1996)] produces a supersensitive psychomotor response in haloperidol-treated rats. For comparison, we also assessed effects of the monoamine reuptake blocker cocaine (Rothman et al., 2001) [0, 2.5 or 10 mg/kg (Kosten, 1997)] and the D1/D2/monoamine receptor agonist apomorphine (Millan et al., 2002) [0, 0.25 or 0.5 mg/kg (Geyer et al., 1987; Barros et al., 1989)]. Each rat was tested 4 times, and received no more than 3 agonist injections (n = 7-14/dose/group) and no more than 2 vehicle injections (vehicle condition: n = 29-31/group).

#### 2.7.2. Experiment 2: Stimulation of D1- or D2-mediated neurotransmission

We determined whether selective D1 or D2 stimulation produces an enhanced psychomotor response in haloperidol-treated rats. Locomotion was recorded for 30 min before administration of a D1 partial agonist [SKF38393 (Seeman and Van Tol, 1994; Neumeyer et al., 2003); 0, 1 or 10 mg/kg (Molloy and Waddington, 1987; Meller et al., 1988)] or a D2 agonist [quinpirole (Millan et al., 2002); 0, 0.15 or 0.5 mg/kg (Benaliouad et al., 2009; Hashimoto et al., 2018)], and for 2 hours thereafter. Each rat received one dose of each agonist (n = 8/dose/group) and one vehicle injection (n = 16/group).

#### 2.7.3. Experiment 3: Blockade of D1- or D2-mediated neurotransmission

We assessed whether D1 and/or D2 transmission is necessary for the expression of dopamine supersensitivity. Rats received the D2 antagonist sulpiride (Caley and Weber, 1995; Martelle and Nader, 2008) [0, 25 or 80 mg/kg (Fritts et al., 1997; Wright et al., 2013)] or the D1 antagonist SCH39166 (McQuade et al., 1991) [0, 0.03 or 0.1 mg/kg (Batsche et al., 1994; Scardochio and Clarke, 2013)], and 30 min later, they received d-amphetamine (0 or 1.5 mg/kg). Rats received 4 out of 12 combinations: 1) vehicle + vehicle (n = 31-32/group), 2) antagonist + vehicle (n = 7-8/combination/group), 3) vehicle + d-amphetamine (n = 31-32/group) or 4) antagonist + d-amphetamine (n = 7-8/combination/group).

#### 2.7.4. Experiment 4: D-amphetamine effects on D1- and D2-mediated signalling in the striatum

Experiments 2-3 showed that both D1- and D2-mediated signalling modulate the supersensitive response to d-amphetamine in antipsychotic-treated rats. Here, we measured d-amphetamine-induced changes in the activity of dopamine receptor-dependent intracellular proteins in the striatum. Specifically, we quantified d-amphetamine-induced protein activity in AKT/GSK3β- and cAMP/PKA-dependent signalling pathways. Locomotion was recorded for 30 minutes, then control and haloperidol-treated rats received s.c. saline (n = 6/group) or d-amphetamine (n = 6/group). One hour later, brains were extracted, samples were taken from the nucleus accumbens and the dorsal, ventrolateral and centromedial caudate-putamen. We quantified total and phosphorylated protein levels of DARPP-32, ERK1, ERK2, AKT and GSK3β using Western Blot procedures.

#### 2.7.5. Experiment 5: Intra-VTA infusion of neurotensin or DAMGO

The preceding experiments suggested that antipsychotic-evoked supersensitivity is linked to increased dopamine-mediated transmission. Here we determined whether increasing VTA dopamine impulse flow evokes a supersensitive psychomotor response in haloperidol-treated rats. To this end, we evaluated the locomotor response to bilateral intra-VTA infusions of vehicle, neurotensin (1 nmol/hemisphere) or DAMGO [a µ-opioid receptor agonist (Chen et al., 1993); 0.3 nmol/hemisphere], at concentrations that increase dopamine release in terminal regions (Kalivas and Duffy, 1990; Laitinen et al., 1990). Neurotensin increases dopamine impulse flow by producing an inward current on dopamine neurons (Mercuri et al., 1993), reducing D2 autoreceptor-mediated inhibition (Werkman et al., 2000; Jomphe et al., 2006; Thibault et al., 2011), and enhancing glutamatergic inputs onto dopamine neurons (Kempadoo et al., 2013; Bose et al., 2015). DAMGO inhibits GABA release, thereby disinhibiting dopamine neuron activity (Kalivas and Duffy, 1990; Bergevin et al., 2002). Here, all rats received vehicle, neurotensin and DAMGO (n = 7/injection/group).

#### 2.7.6. Experiment 6: Intracerebroventricular d-amphetamine

Systemic d-amphetamine administration reliably triggers a supersensitive response in haloperidol-treated rats (Figs. 2A-F) [see also (Smith and Davis, 1975; Ericson et al., 1996; Samaha et al., 2007)], here we determined if limiting d-amphetamine’s effects to the brain is sufficient to produce this effect. We infused d-amphetamine bilaterally into the lateral ventricles [0, 50 or 150 µg/hemisphere (Lin et al., 1983)] and measured psychomotor activity. All rats received vehicle (n = 16-21/group) and one of 2 d-amphetamine concentrations (n = 6-10/concentration/group).

**Fig. 2.**
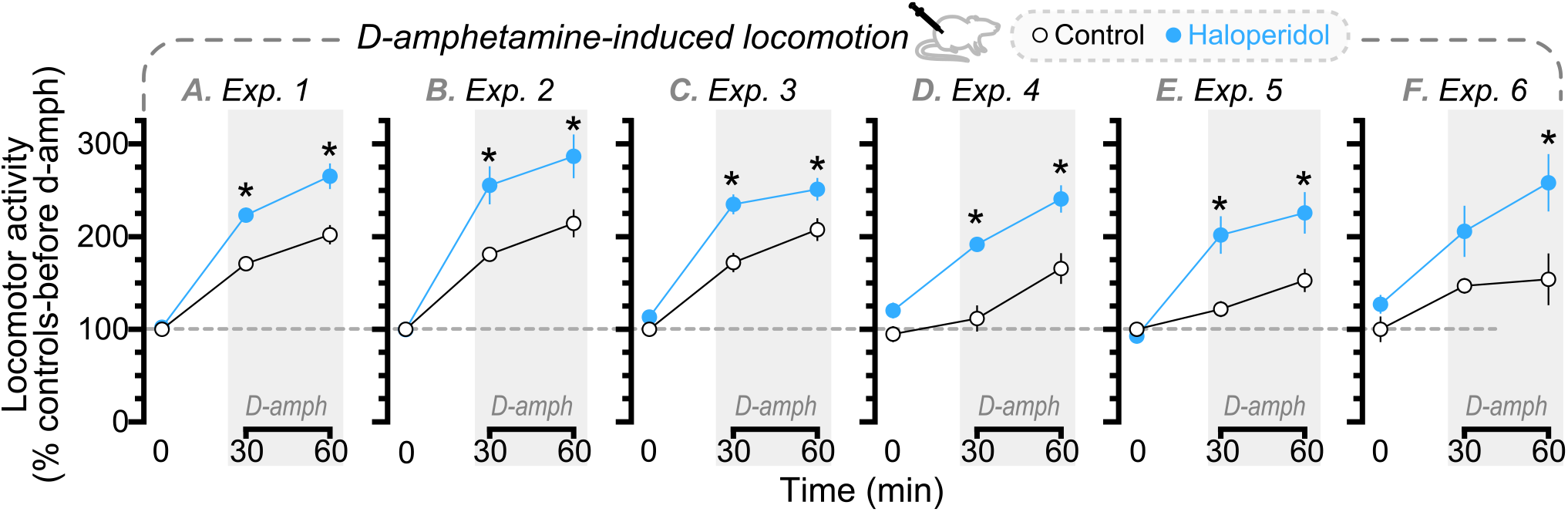
Across experiments, following haloperidol treatment cessation, rats showed enhanced d-amphetamine-induced locomotor activity. (A-F) Locomotor response to subcutaneous (s.c.) d-amphetamine (1.5 mg/kg). In all data panels, dotted lines indicate locomotion of vehicle-injected controls. *n*’s = 5-32/condition. \**p* < 0.05, relative to controls at same time point.

### 2.8. Statistical analysis

In Fig. 2, d-amphetamine-induced locomotion was expressed as the percent change relative to the locomotor activity of controls in the first 30 min of the test session (*i*.*e*., prior to d-amphetamine injection). This allows us to compare the magnitude of the d-amphetamine response relative to baseline levels of locomotor activity under our test conditions. In Figs. 3-7 and Figs. S1-5, data are expressed as the percent change relative to vehicle-injected controls, as the corresponding experiments included a vehicle-injected, control condition. Mixed-model ANOVA was used to analyse the influence of Injection or Group on locomotion (Group × Injection × Time), psychomotor activity ratings or protein level (Group × Injection; ‘Injection’ as a between-subjects variable in Experiments 1-4 and 6, and a within-subjects variable in Experiment 5). When interaction and/or main effects were significant (*p* ≤ 0.05), effects were analysed further using Bonferroni-adjusted multiple post-hoc comparisons. Values in figures are mean ± SEM.

**Fig. 3.**
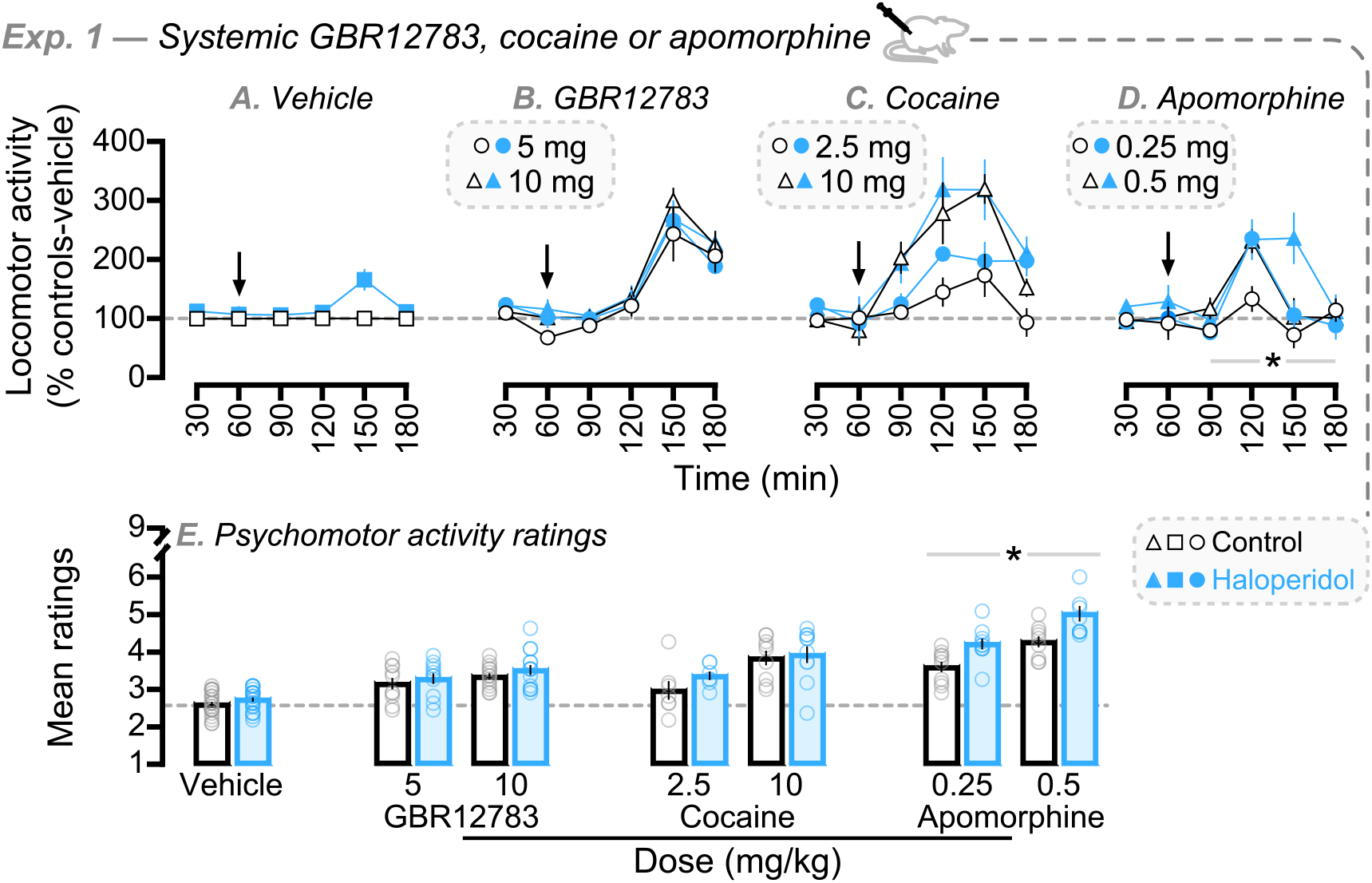
Following discontinuation of chronic haloperidol treatment, rats are supersensitive to the psychomotor activating effects of apomorphine, but not GBR12783 or cocaine. Psychomotor response to (A) vehicle, (B) subcutaneous GBR12783 (5 or 10 mg/kg), (C) intraperitoneal cocaine (2.5 or 10 mg/kg) and (D) subcutaneous apomorphine (0.25 or 0.5 mg/kg). (E) Psychomotor activity ratings after vehicle, GBR12783, cocaine or apomorphine administration. In all data panels, dotted lines indicate response of vehicle-injected controls. *n*’s = 7-31/condition. \**p* < 0.05. In (D); *Group × Injection × Time interaction, Group × Time interaction and Group effects. In (E); *Group effect.

## 3. Results

Across experiments, locomotor activity did not differ between haloperidol-treated and control groups prior to d-amphetamine injection (all *P*’s > 0.05). In Experiment 4, control and haloperidol-treated rats that received vehicle prior to brain extraction also showed similar levels of locomotion (data not shown; *p* > 0.05).

Across experiments, all haloperidol-treated groups developed dopamine supersensitivity, as indicated by enhanced d-amphetamine-induced locomotion relative to controls following antipsychotic treatment cessation (Fig. 2; Group × Time interaction; 2A, *F*_2,102_ = 10.97, *p* < 0.0001; 2B, *F*_2,60_ = 7.74, *p* = 0.001; 2C, *F*_2,122_ = 8.11, *p* = 0.0005; 2D, *F*_2,20_ = 4.55, *p* = 0.024; 2E, *F*_2,22_ = 18.31, *p* < 0.0001; 2F, *F*_2,16_ = 3.83, *p* = 0.044; Group effect; 2A, *F*_1,51_ = 15.76, *p* = 0.0002; 2B, *F*_1,30_ = 8.4, *p* = 0.007; 2C, *F*_1, 61_ = 11.17, *p* = 0.001; 2D, *F*_1,10_ = 23.15, *p* = 0.0007; 2E, *F*_1,11_ = 7.92, *p* = 0.017; 2F, *F*_1,8_ = 5.74, *p* = 0.043; haloperidol > controls; 2A; min 30, *p* = 0.0002; min 60, *p* < 0.0001; 2B; min 30, *p* = 0.0018; min 60, *p* = 0.0026; 2C; min 30, *p* < 0.0001; min 60, *p* = 0.006; 2D; min 30, *p* = 0.0002; min 60, *p* = 0.0004; 2E; min 30, *p* = 0.0008; min 60, *p* = 0.002; 2F; min 60, *p* = 0.008). D-amphetamine increased psychomotor activity ratings beyond vehicle, without evidence of stereotypy (i.e., mean score < 6), and this was similar across groups (Fig. S1).

### 3.1. Experiment 1: Stimulation of dopamine neurotransmission

GBR12783, cocaine and apomorphine increased locomotion above vehicle (Figs. 3A-D; minutes 90-180; Injection × Time interaction; 3A versus 3B, *F*_6,309_ = 14.6, *p* < 0.0001; 3A versus 3C, *F*_6,261_ = 6.28, *p* < 0.0001; 3A versus 3D, *F*_6,279_ = 8.61, *p* < 0.0001; Injection effect; 3A versus 3B, *F*_2,103_ = 31.78, *p* < 0.0001; 3A versus 3C, *F*_2,87_ = 46.63, *p* < 0.0001; 3A versus 3D, *F*_2,93_ = 9.8, *p* < 0.0001). There were no group differences in GBR12783- or cocaine-induced locomotion (Figs. 3B-C; all *P’s* > 0.05). However, haloperidol rats showed greater apomorphine-induced locomotion relative to controls (Fig. 3D; Group × Injection × Time interaction, *F*_3,105_ = 3.03, *p* = 0.033; Group × Time interaction, *F*_3,105_ = 3.68, *p* = 0.014; Group effect, *F*_1,35_ = 5.18, *p* = 0.029) [see also (Smith and Davis, 1975; Montanaro et al., 1982; Carvalho et al., 2009)].

Similarly, GBR12783, cocaine and apomorphine increased psychomotor activity ratings compared to vehicle (Fig. 3E; Injection effect; vehicle versus GBR12783, *F*_2,91_ = 41.4, *p* < 0.0001; vehicle versus cocaine, *F*_2,75_ = 53.45, *p* < 0.0001; vehicle versus apomorphine, *F*_2,81_ = 189.8, *p* < 0.0001), and haloperidol rats had greater ratings compared to controls in response to apomorphine (Fig. 3E; apomorphine; Group effect, *F*_1,35_ = 17.73, *p* = 0.0002). No other comparisons were statistically significant.

Thus, following cessation of haloperidol treatment, rats with antipsychotic-induced supersensitivity showed a supersensitive psychomotor response to a monoamine receptor agonist (apomorphine), but not to a monoamine reuptake blocker (cocaine) or a selective dopamine reuptake inhibitor (GBR12783).

### 3.2. Experiment 2: Stimulation of D1- or D2-mediated neurotransmission

Across groups, the D2 agonist quinpirole dose-dependently increased locomotion relative to vehicle (Figs. 4A versus 4B; minutes 60-210; Injection × Time interaction, *F*_10,290_ = 26.66, *p* < 0.0001; Injection effect, *F*_2,58_ = 32.51, *p* < 0.0001), and this was greatest in haloperidol rats (Fig.4B; minutes 60-210; Group × Time interaction, *F*_5,140_ = 4.16, *p* = 0.001; Group effect, *F*_1,28_ = 3.93, *p* = 0.057). The D1 partial agonist SKF38393 did not increase locomotion (Figs. 4A versus 4C), consistent with findings that it evokes stereotypy, but little hyperlocomotion (Meller et al., 1988; Meyer and Shults, 1993; Hooks et al., 1994). Indeed, both quinpirole and SKF38393 increased psychomotor activity ratings relative to vehicle (Fig. 4D; Injection effect; vehicle versus quinpirole, *F*_2,58_ = 17.85, *p* < 0.0001; vehicle versus SKF38393, *F*_2,58_ = 15.89, *p* < 0.0001), with the highest dose of each agonist producing greater effects (Fig. 4D; Injection effect; quinpirole, *F*_1,28_ = 12.14, *p* = 0.0016; SKF38393, *F*_1,28_ = 9.41, *p* = 0.0048). There were no group differences in these effects.

**Fig. 4.**
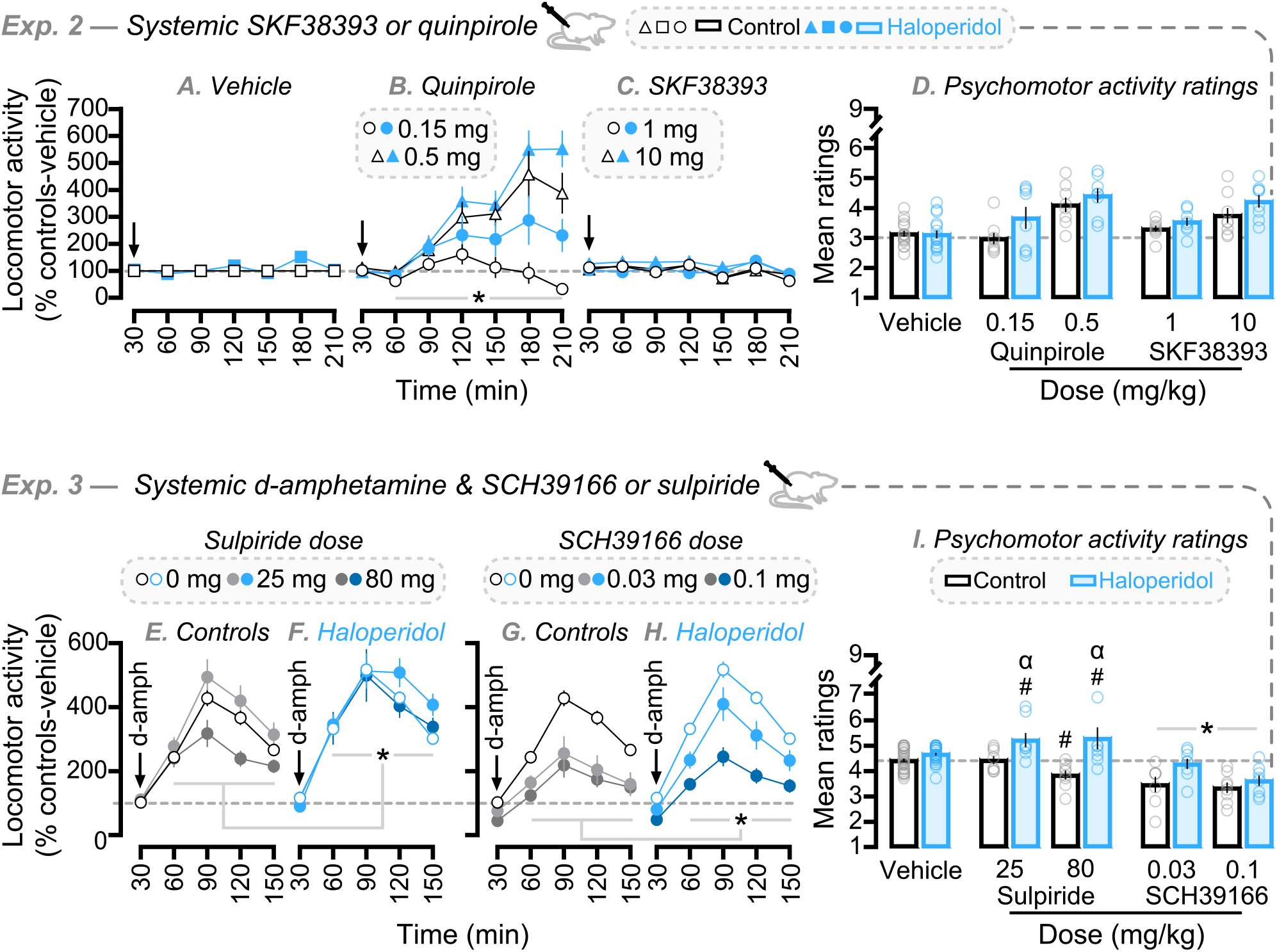
Both D1 receptor- and D2 receptor-mediated neurotransmission regulate the expression of antipsychotic-evoked dopamine supersensitivity. Locomotor response to subcutaneous (A) vehicle, (B) quinpirole (0.15 or 0.5 mg/kg) or (C) SKF38393 (1 or 10 mg/kg). (D) Psychomotor activity ratings after vehicle, quinpirole or SKF38393 administration. Effects of subcutaneous (E-F) sulpiride (0, 25 or 80 mg/kg) or (G-H) SCH39166 (0, 0.03 or 0.1 mg/kg) on the locomotor response to subcutaneous d-amphetamine. (I) Effects of sulpiride or SCH39166 on d-amphetamine-induced psychomotor activity ratings. In all data panels, dotted lines indicate response of vehicle-injected controls. *n*’s = 7-32/condition. ^α/#/^\**p* < 0.05. In (B); *Group × Time interaction and Group effects. In (E-F) and (G-H); *Group effect. In (I); ^#^versus vehicle from same group, ^α^versus controls at same sulpiride dose, *Group effect.

Thus, rats that express dopamine supersensitivity following discontinuation of chronic antipsychotic exposure showed an augmented behavioural response to D2, but not D1 receptor stimulation.

### 3.3. Experiment 3: Blockade of D1- or D2-mediated neurotransmission

Across groups, the D2 antagonist sulpiride did not influence vehicle-induced locomotion or ratings (Figs. S2A-B and S2E). However, sulpiride influenced d-amphetamine-induced locomotion (Figs. 4E-F; minutes 60-150; Injection × Time interaction, *F*_6,267_ = 3.11, *p* = 0.006; Injection effect, *F*_2,89_ = 3.06, *p* = 0.052), and did so in a group-dependent manner (Figs. 4E-F; Group effect, *F*_1,89_ = 13.64; *p* = 0.0004). In fact, the ratings showed that sulpiride had opposing effects on d-amphetamine-induced psychomotor activity in controls and haloperidol-treated rats. Indeed, sulpiride *suppressed* d-amphetamine-induced psychomotor activity ratings in controls but, surprisingly, it *enhanced* this response in haloperidol-treated rats (Fig. 4I; vehicle and sulpiride; Group × Injection interaction, *F*_2,89_ = 8.47, *p* = 0.0004; Group effect, *F*_1,89_ = 38.25, *p* < 0.0001; haloperidol > controls; 25 mg/kg, *p* = 0.012; 80 mg/kg, *p* < 0.0001; haloperidol rats; 0 < 25 mg/kg, *p* = 0.031; 0 < 80 mg/kg, *p* = 0.013; control rats; 0 > 80 mg/kg, *p* = 0.024).

The D1 antagonist SCH39166 reduced vehicle-induced locomotion and ratings across groups (Figs. S2C-E). Haloperidol-treated rats showed greater d-amphetamine-induced locomotion and ratings than controls, regardless of SCH39166 injection (Figs. 4G-H; minutes 60-150; Group effect, *F*_1,88_ = 6.92, *p* = 0.01; Fig. 4I; Group effect across vehicle and SCH39166 ratings, *F*_1,88_ = 14.48, *p* = 0.0003). SCH39166 also decreased d-amphetamine-induced locomotion and ratings across groups (Figs. 4G-H; minutes 60-150; Injection × Time interaction, *F*_6,264_ = 6.15, *p* < 0.0001; Injection effect, *F*_2,88_ = 28.25, *p* < 0.0001; Fig. 4I; Injection effect across vehicle and SCH39166 ratings, *F*_2,88_ = 39.11, *p* < 0.0001). Notably, in haloperidol-treated rats, 0.03 mg/kg SCH39166 restored d-amphetamine-induced locomotion to control levels (compare light blue curve in Fig. 4H to white curve in Fig. 4G).

Thus, in rats that had previously received an antipsychotic treatment producing dopamine supersensitivity, blockade of D1 receptors *tempered*, while blockade of D2 receptors *potentiated* the enhanced psychomotor response to d-amphetamine.

### 3.4. Experiment 4: D-amphetamine effects on D1- and D2-mediated signalling in the striatum

#### 3.4.1. Caudate-putamen

D-amphetamine produced a similar signalling profile in haloperidol-treated and control rats. Across groups, d-amphetamine did not increase protein phosphorylation in AKT/GSK3β- or cAMP/PKA-dependent pathways in the caudate-putamen (Figs. S3-5).

#### 3.4.2. Nucleus accumbens

See Fig. S6-7 in Supplement for full pictures of Western blots shown in Fig. 5. Relative to saline, d-amphetamine increased total GSK3β levels only in haloperidol-treated rats (Fig. 5A; Group × Injection interaction, *F*_1,20_ = 4.23, *p* = 0.053; Injection effect, *F*_1,20_ = 14.61, *p* = 0.011; haloperidol rats; d-amphetamine > saline, *p* = 0.001). This reflects higher levels of non-phosphorylated (active) versus phosphorylated (inactive) GSK3β (Sutherland et al., 1993), because d-amphetamine decreased pGSK3β/total GSK3β ratios across groups (Fig. 5B; Injection effect, *F*_1,20_ = 7.57, *p* = 0.012). D-amphetamine decreased total AKT levels and increased pAKT/total AKT ratios, with no group differences (Injection effect; Fig. 5C; *F*_1,15_ = 13.01, *p* = 0.0026; Fig. 5D; *F*_1,15_ = 6.61, *p* = 0.021). Hence, in the nucleus accumbens, dopamine-supersensitive and control rats show similar d-amphetamine-induced effects on AKT, but dopamine-supersensitive rats show greater d-amphetamine-induced increases in GSK3β activity.

**Fig. 5.**
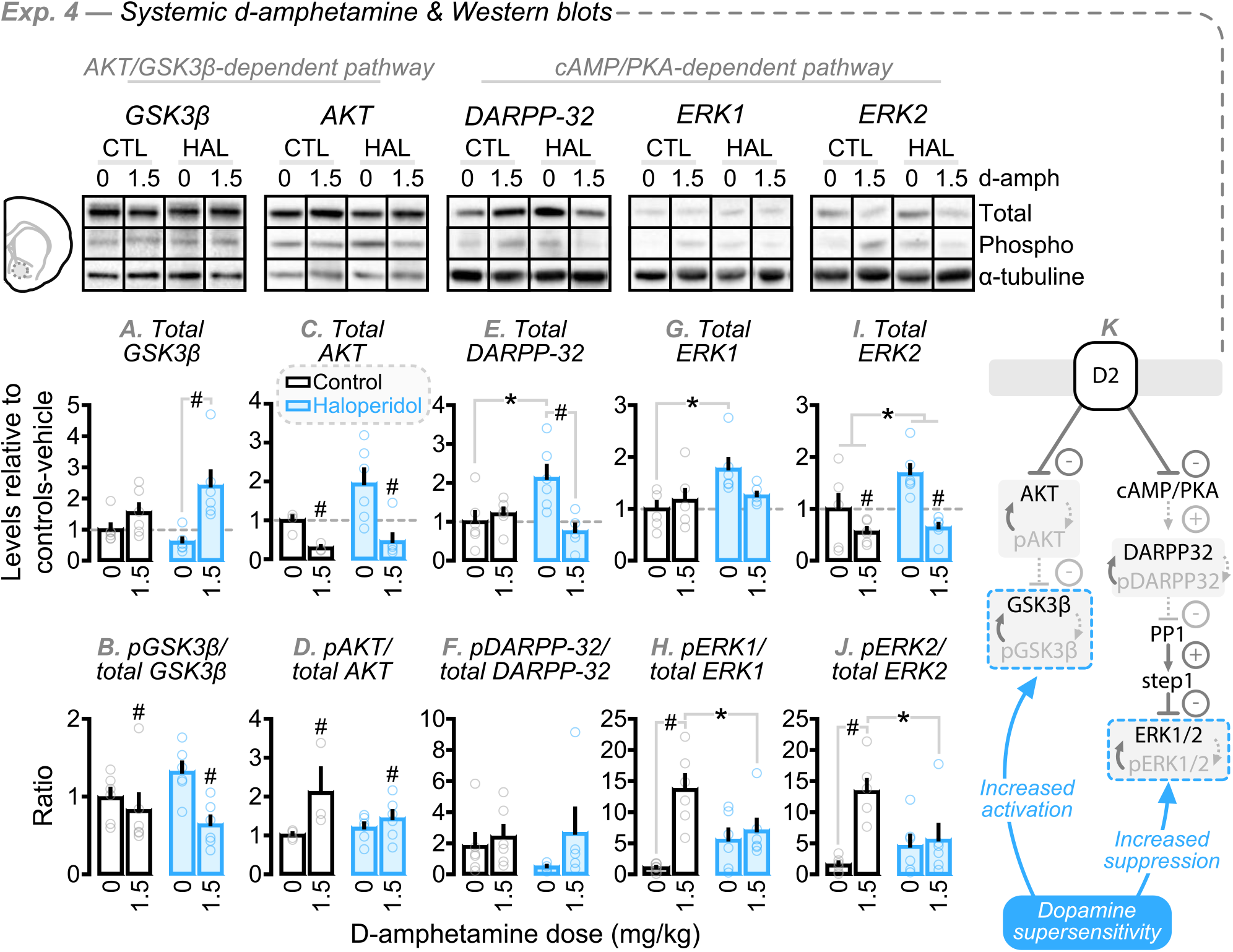
Rats with antipsychotic-induced dopamine-supersensitivity have enhanced d-amphetamine-induced GSK3β activity and suppressed d-amphetamine-induced ERK1/2 activity in the nucleus accumbens following antipsychotic treatment cessation. Top illustration, Western blots in accumbens tissue. Total protein levels and phosphorylated/total protein ratios within the (A-D) AKT/GSK3β- and (E-J) cAMP/PKA-dependent pathways. In (A, C, E, G, I), dotted lines indicate mean protein level of vehicle-injected controls. (K) Proposed model of d-amphetamine’s D2 receptor-mediated biochemical effects in rats with antipsychotic-evoked dopamine supersensitivity. *n*’s = 3-6/condition. \**p* < 0.05; in (I), Group effect. ^#^*p* < 0.05; in (B-D, I), Injection effect.

In haloperidol-treated rats, total DARPP-32 levels were increased at baseline and decreased after d-amphetamine (Fig. 5E; Group × Injection interaction, *F*_1,19_ = 9.97, *p* = 0.005; Injection effect, *F*_1,19_= 5.47, *p* = 0.03; after saline; haloperidol rats > controls, *p* = 0.0086; Haloperidol rats, saline > d-amphetamine, *p* = 0.0024). At baseline, total levels of both ERK1 and ERK2 were highest in haloperidol-treated rats (Fig. 5G; Group × Injection interaction, *F*_1,20_ = 4.13, *p* = 0.056; Group effect, *F*_1,20_ = 6.41, *p* = 0.02; haloperidol rats > controls after saline injection, *p* = 0.0084; Fig. 5I; Group effect, *F*_1,20_ = 4.56, *p* = 0.045). D-amphetamine decreased total ERK2 levels similarly across groups (Fig. 5I; Injection effect, *F*_1,20_ = 17.79, *p* = 0.0004). Hence, d-amphetamine-induced expression of dopamine supersensitivity potentially involves decreased total DARPP-32 levels in the accumbens, without distinct effects on total ERK1/ERK2 levels.

D-amphetamine enhanced the proportion of phosphorylated (active) versus total ERK1 and ERK2 levels in controls [also see (Svenningsson et al., 2003; Valjent et al., 2005; Beaulieu et al., 2006)], but not in haloperidol-treated rats (Fig. 5H; Group × Injection interaction, *F*_1,20_ = 9.73, *p* = 0.0054; Injection effect, *F*_1,20_ = 15.62, *p* = 0.0008; Fig. 5J; Group × Injection interaction, *F*_1,20_ = 8.41, *p* = 0.009; Injection effect, *F*_1,20_ = 11.82, *p* = 0.0026; controls; d-amphetamine > saline; Fig. 5H, *p* = 0.0001; Fig. 5J, *p* = 0.0005; after d-amphetamine; controls > haloperidol rats; Fig. 5H, *p* = 0.031; Fig. 5J, *p* = 0.016). D-amphetamine did not change the proportion of phosphorylated DARPP-32 in either group (Fig. 5F; *p* > 0.05). Thus, in the accumbens, the expression of dopamine supersensitivity is potentially linked to suppressed phosphorylation of ERK1 and ERK2.

Thus, d-amphetamine does not produce consistent effects in the caudate-putamen of rats with established antipsychotic-induced dopamine supersensitivity. However, enhanced d-amphetamine-induced psychomotor activity in these rats was accompanied by increased GSK3β activity and decreased ERK activity in the nucleus accumbens, which could reflect greater D2-mediated transmission (Fig. 5K).

### 3.5. Experiment 5: Intra-VTA infusion of neurotensin or DAMGO

Intra-VTA neurotensin enhanced locomotion beyond vehicle (Figs. 6B versus 6C; minutes 30-120; Injection × Time interaction, *F*_3,36_ = 8.68, *p* = 0.0002; Injection effect, *F*_1,12_ = 13.77, *p* = 0.003). There was a significant Injection × Group × Time interaction effect (Figs. 6B-C; minutes 30-120; *F*_3,36_ = 3.18, *p* = 0.036), but further analyses show that controls and haloperidol-treated rats had a similar locomotor response to vehicle (Fig. 6B, *p* > 0.05) and neurotensin (Fig. 6C; *p* > 0.05). Intra-VTA DAMGO increased locomotion beyond vehicle (Figs. 6B versus 6D; minutes 30-120; Injection × Time interaction, *F*_3,36_ = 12.08, *p* < 0.0001; Injection effect, *F*_1,12_ = 17.46, *p* = 0.001), without group differences (Fig. 6D; *p* > 0.05). Both neurotensin and DAMGO increased ratings above vehicle, and did so similarly across groups (Fig. 6E; Injection effect; vehicle versus neurotensin, *F*_1,12_ = 6.92, *p* = 0.022; vehicle versus DAMGO, *F*_1,12_ = 13.91, *p* = 0.0029).

**Fig. 6.**
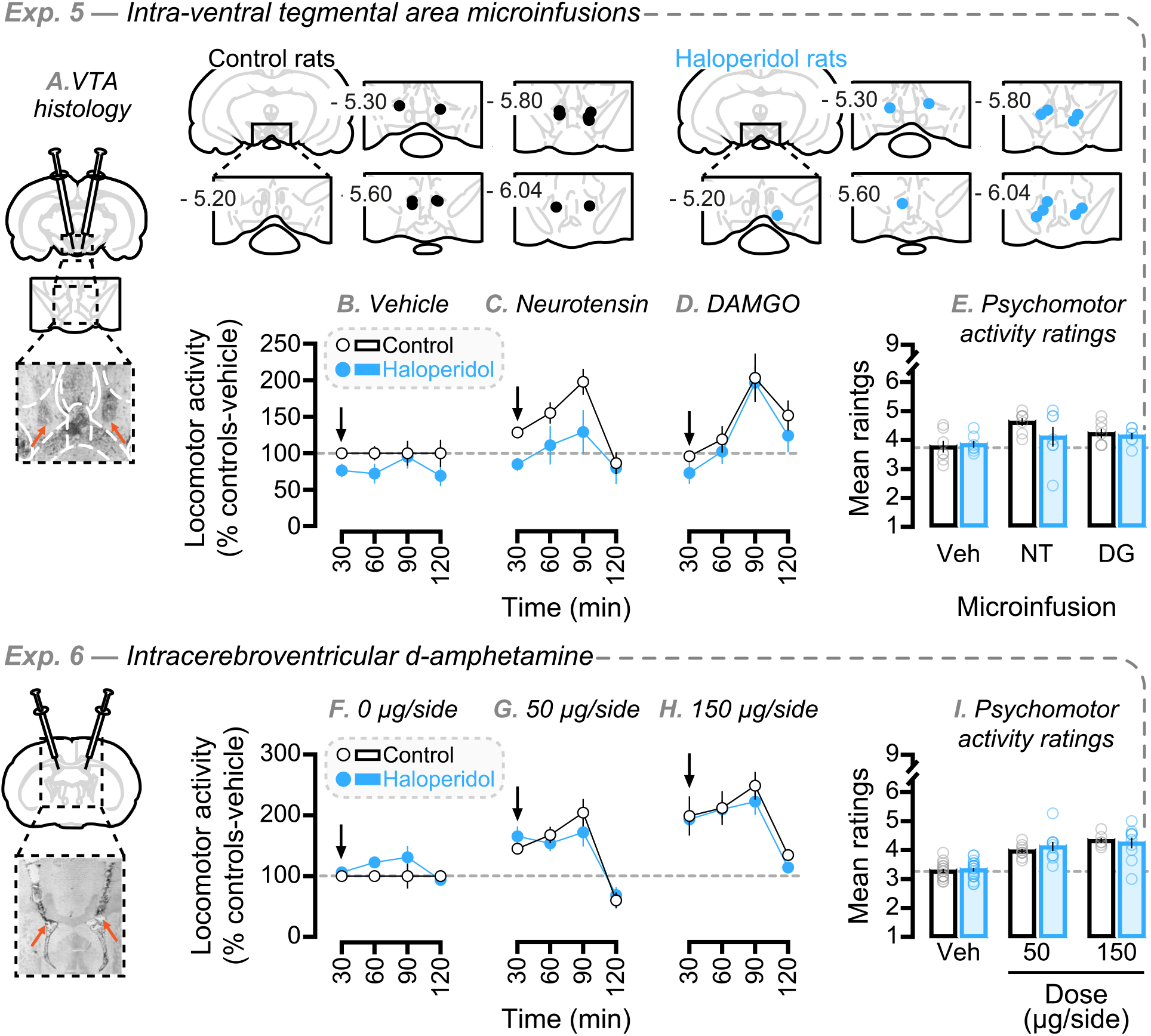
Neither increasing ventral tegmental area (VTA) dopamine impulse flow nor injecting d-amphetamine into the lateral ventricles triggers the expression of established antipsychotic-evoked dopamine supersensitivity. (A) VTA histology. (B-D) Locomotor response to intra-VTA vehicle, neurotensin (1 nmol/hemisphere) or DAMGO (0.3 nmol/hemisphere). (E) Psychomotor activity ratings following intra-VTA vehicle, neurotensin (NT) or DAMGO (DG). (F-H) Locomotor response to intracerebroventricular d-amphetamine (0, 50 or 150 µg/hemisphere). (I) Psychomotor activity ratings evoked by intracerebroventricular d-amphetamine. Bottom left, representative injector placements (orange arrows indicate injectors). In all data panels, dotted lines indicate response of vehicle-injected controls. *n*’s = 7-21/condition.

Thus, while increasing VTA dopamine impulse flow enhanced psychomotor activity in both controls and rats that had previously received chronic haloperidol treatment, it did not trigger a supersensitive response in the haloperidol-treated group.

### 3.6. Experiment 6: Intracerebroventricular d-amphetamine

Across groups, intracerebroventricular d-amphetamine dose-dependently increased locomotion and psychomotor activity ratings compared to vehicle (Figs. 6F versus 6G-H; minutes 30-120; Injection × Time interaction, *F*_6,198_ = 9,86, *p* < 0.0001; Injection effect, *F*_2,66_ = 29.23, *p* < 0.0001; Fig. 6I; Injection effect, *F*_2,66_ = 50.43, *p* < 0.0001), without group differences (Figs. 6G-I; all *P*’s > 0.05). This indicates that injecting d-amphetamine into the cerebral ventricles produced control levels of psychomotor activity in rats previously treated with haloperidol.

Thus, injecting neurotensin or DAMGO into the VTA or d-amphetamine into the lateral ventricles does not trigger the behavioural expression of antipsychotic-evoked dopamine supersensitivity (Fig. 6). One possible explanation for this is that these manipulations produced ceiling levels of locomotor activity, such that dopamine-supersensitive rats could not show any further increases in locomotion. To address this possibility, we compared the degree of psychomotor activity produced by s.c. d-amphetamine, intra-VTA neurotensin or DAMGO (Figs. 7A-B), and i.c.v. d-amphetamine (Figs. 7C-D). This analysis suggests that the intracranial treatments did not produce maximal levels of locomotion in antipsychotic-treated rats. Specifically, in these rats, the magnitude of the psychomotor response to s.c. d-amphetamine was significantly higher compared to that evoked by the intracranial treatments. As Figs. 7A-B show, in Experiment 5, s.c. d-amphetamine produced a greater locomotor response relative to intra-VTA neurotensin or DAMGO in haloperidol-treated rats (but not in controls; Figs. 7A-B; Group × Injection interaction, *F*_2,35_ = 7.43, *p* = 0.0021; Injection effect, *F*_2,35_ = 6.22, *p* = 0.0049; haloperidol-treated group: s.c. d-amphetamine > intra-VTA neurotensin, *p* < 0.0001; s.c. d-amphetamine > intra-VTA DAMGO, *p* = 0.0013). Similarly, Figs. 7C-D also show that in Experiment 6, s.c. d-amphetamine produced a greater psychomotor response relative to i.c.v. d-amphetamine (50 µg/side) in haloperidol-treated rats (but not in control rats; Figs. 7C-D; Group × Injection interaction, F_2,39_ = 4.37, *p* = 0.02; Injection effect, *F*_2,39_ = 7.33, *p* = 0.002; in haloperidol-treated group: s.c. d-amphetamine > 50 µg/side d-amphetamine, *p* = 0.0009).

**Fig. 7.**
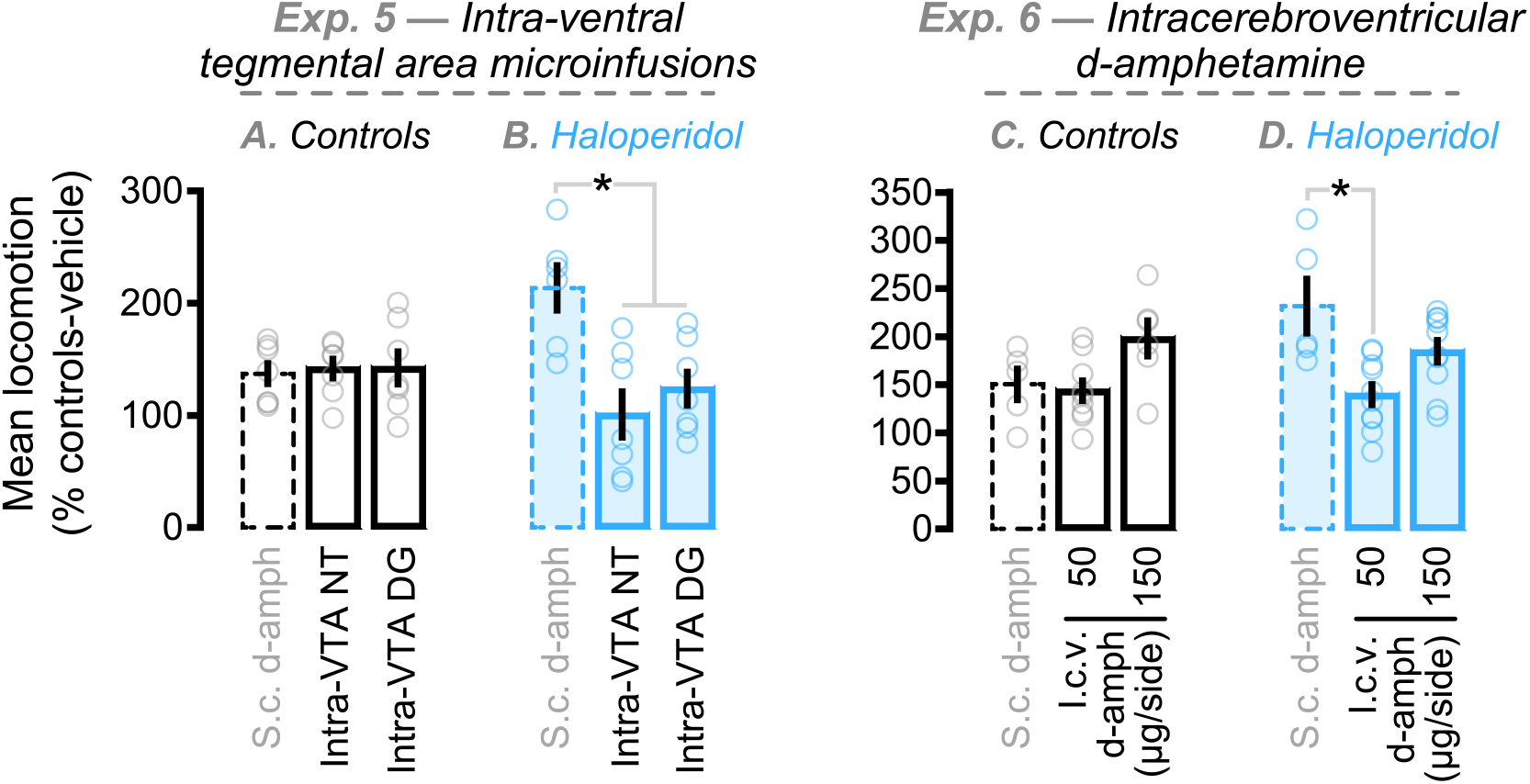
Degree of psychomotor activity produced by subcutaneous (s.c.) d-amphetamine, intra-ventral tegmental area (VTA) neurotensin, intra-VTA DAMGO or intracerebroventricular (i.c.v) d-amphetamine. In Experiment 5, (**A**) controls and (**B**) haloperidol-treated rats received s.c. d-amphetamine, intra-VTA neurotensin or DAMGO. In Experiment 6, (**C**) controls and (**D**) haloperidol-treated rats received s.c. or i.c.v. d-amphetamine (50 or 150 µg/side). Values are mean locomotor activity relative to vehicle-injected controls. *n*’s = 5-10/condition. \**p* < 0.05.

Another possibility is that injecting d-amphetamine into the cerebral ventricles does not trigger an exaggerated psychomotor response in haloperidol-pretreated rats, because i.c.v. d-amphetamine diffuses to fewer sites of action compared to s.c. d-amphetamine. However, the findings suggest that i.c.v. d-amphetamine can be as potent as s.c. d-amphetamine in enhancing psychomotor activity. Indeed, the level of psychomotor activity evoked by i.c.v. versus s.c. d-amphetamine was similar in both control rats (s.c. d-amphetamine vs 50 and 150 µg/side, all *P*’s > 0.05) and in haloperidol-treated rats (s.c. d-amphetamine vs relative to 150 µg/side, *p* > 0.05).

## 4. Discussion

Rats received a clinically relevant haloperidol treatment regimen, and they developed dopamine supersensitivity, as indicated by an enhanced psychomotor response to systemic d-amphetamine after antipsychotic treatment cessation (Smith and Davis, 1975; Ericson et al., 1996; Samaha et al., 2007). We characterized this dopamine supersensitive state evoked by antipsychotics, and we report four key findings. First, dopamine-supersensitive rats showed a normal psychomotor response to both the dopamine reuptake inhibitors GBR12783 and cocaine and to the D1 receptor partial agonist SKF38393. In contrast, they showed an enhanced response to the D2 receptor agonist quinpirole and the non-selective dopamine receptor agonist apomorphine. Thus, direct stimulation of D2, but not D1 receptors triggers the expression of dopamine supersensitivity. Second, D1 and D2 antagonism produced opposite behavioural effects in dopamine-supersensitive rats. Blockade of D2 receptors *enhanced* the exaggerated psychomotor response to d-amphetamine, whereas blockade of D1 receptors *suppressed* it. This indicates that D1, but not D2 receptor activity is necessary for the expression of dopamine supersensitivity, further supporting dissociable roles for D1-versus D2-mediated neurotransmission. Third, the expression of antipsychotic-evoked supersensitivity is linked to changes in GSK3β and ERK activity in the nucleus accumbens (but not caudate-putamen) that are consistent with enhanced D2-mediated transmission. Fourth, pharmacologically increasing VTA dopamine impulse flow or intracerebroventricular d-amphetamine administration both produced a normal psychomotor response in dopamine-supersensitive rats. This suggests that increasing monoaminergic transmission in the brain is not sufficient to trigger the expression of established dopamine supersensitivity. Thus, while both D1- and D2-dependent neurotransmission mediate the expression of dopamine supersensitivity in rats with a history of chronic haloperidol treatment, this supersensitive response requires more than just an increase in extracellular dopamine concentrations and dopamine neurotransmission in the brain (Fig. 8).

**Fig. 8.**
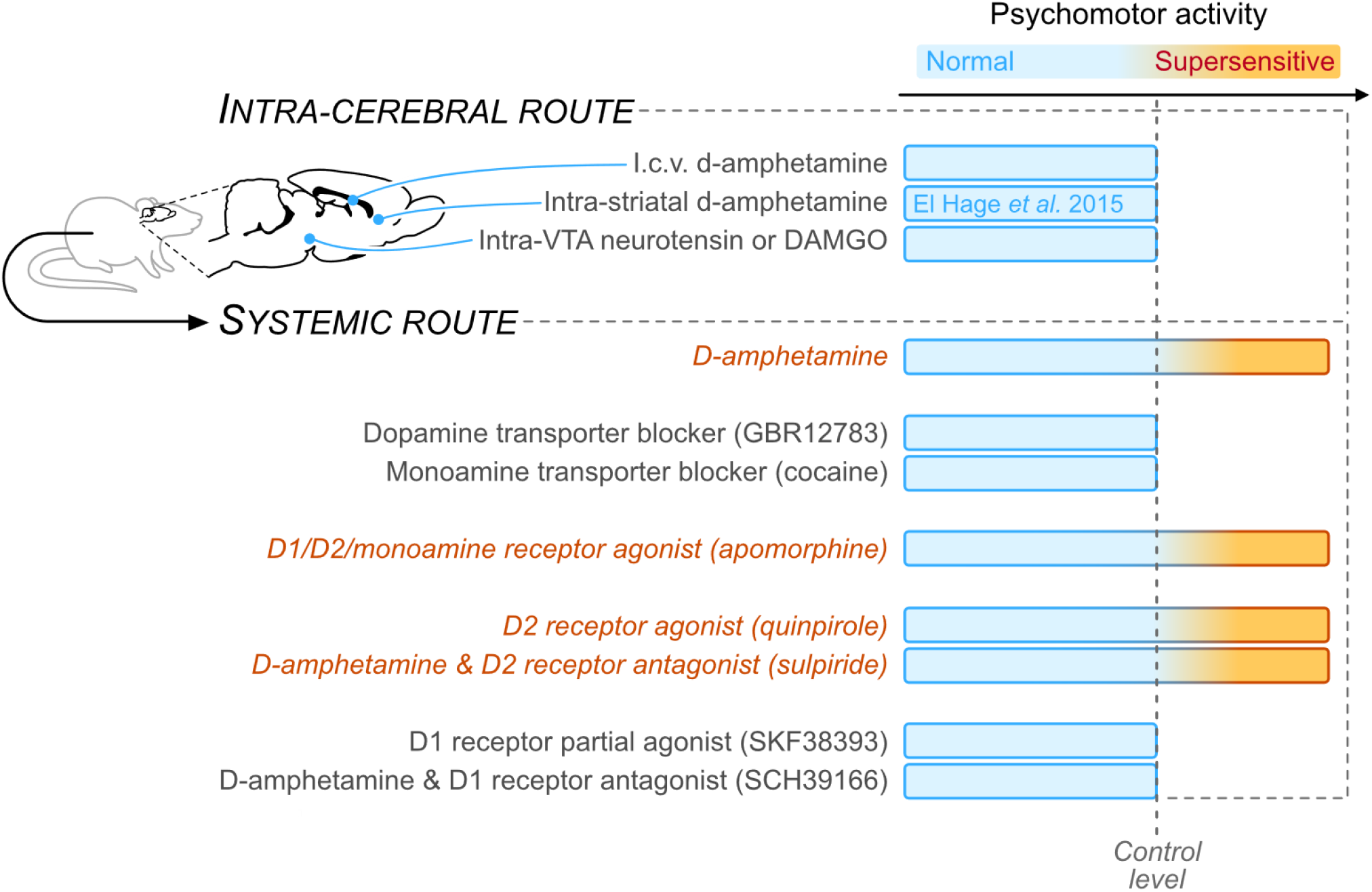
Conceptual summary. Rats exposed to continuous haloperidol show a supersensitive psychomotor response to subcutaneous d-amphetamine. However, the same rats show a normal psychomotor response to intra-striatal and intracerebroventricular d-amphetamine, as well as to intra-ventral tegmental area (VTA) infusions of neurotensin or DAMGO (a manipulation enhancing dopamine impulse flow). Furthermore, when injected through the systemic route, dopamine-supersensitive rats have a greater psychomotor response to the monoamine receptor agonist apomorphine and the D2 receptor agonist quinpirole, while they show normal psychomotor activity in response to the dopamine transporter blocker GBR12783, the monoamine transporter blocker cocaine and the D1 receptor partial agonist SKF38393. Lastly, the D1 receptor antagonist SCH39166 but not the D2 receptor antagonist sulpiride inhibits the exaggerated psychomotor response to d-amphetamine. Note that all effects summarized in this figure stem from work presented in this article, with the exception of the response to intra-striatal d-amphetamine, which was reported in our previous work (El Hage et al., 2015).

### 4.1. D1 and D2 receptor contributions

Our results suggest that the expression of dopamine supersensitivity following discontinuation of antipsychotic treatment involves enhanced D2-mediated activity. First, our dopamine-supersensitive rats showed an enhanced psychomotor response to a D2 receptor agonist. This could involve the ability of chronic antipsychotic treatment to increase striatal D2 density and function (Burt et al., 1977; Clow et al., 1980; Fleminger et al., 1983; Samaha et al., 2007; Samaha et al., 2008). Second, acute injection of the D2 receptor antagonist, sulpiride suppressed the psychomotor response to d-amphetamine in controls (80 mg/kg sulpiride), but it *potentiated* d-amphetamine responding in dopamine-supersensitive rats (25 and 80 mg/kg sulpiride). This is reminiscent of earlier findings showing that some D2 antagonists—including sulpiride—can potentiate the psychomotor effects of d-amphetamine (Schaefer and Michael, 1984; Robertson and MacDonald, 1985), but this effect is reported at lower sulpiride doses than used here. It is possible that higher sulpiride doses than used here might suppress d-amphetamine-induced locomotion in dopamine-supersensitive rats. However, such a finding would not change the conclusion that these rats are tolerant to the antidopaminergic effects of sulpiride, because our findings already show that a dose of sulpiride (80 mg/kg) that suppresses amphetamine-induced locomotion in control rats has no such suppressive effects in rats previously exposed to chronic haloperidol (Figs. 4E-F). Moreover, it is not clear why sulpiride potentiated d-amphetamine-induced locomotion in dopamine-supersensitive rats, but not in controls. This could involve sulpiride interactions with residual haloperidol at the D2 receptors. However, this is unlikely. In rats, striatal D2 receptor occupancy declines markedly 24 hours after haloperidol administration (Kapur et al., 2003), and here we tested the rats at least 3 days after the cessation of haloperidol treatment. It is also possible that sulpiride potentiates the behavioural response to d-amphetamine in dopamine-supersensitive rats in part by inhibiting presynaptic, D2 autoreceptors, thereby promoting dopamine release and locomotor activity. If this is true, it would suggest that antipsychotic treatment regimens that produce dopamine supersensitivity alter D2 autoreceptor function in ways that enhance the ability of antagonists like sulpiride to promote presynaptic dopamine release. In support, chronic antipsychotic treatment can enhance presynaptic, D2 autoreceptor activity in the caudate-putamen (Calabresi et al., 1992) [but not in the nucleus accumbens (Chesi et al., 1995)].

Finally, dopamine-supersensitive rats showed d-amphetamine-induced changes in nucleus accumbens cAMP/PKA- and GSK3β/AKT-dependent activity consistent with enhanced D2-mediated signalling (Fig. 5K). D2 receptor stimulation enhances GSK3β activity and supresses ERK1/2 phosphorylation (Cross et al., 1995; Nishi et al., 1997; Beaulieu et al., 2007; Oda et al., 2015). Here, d-mamphetamine administration potentiated these biochemical responses in the accumbens of antipsychotic-treated rats. Our biochemical and behavioural findings are correlational, and as discussed below, the central effects of d-amphetamine are likely not sufficient to uncover antipsychotic-evoked dopamine supersensitivity. However, it remains to be determined if the observed changes in GSK3β/ERK activity are *necessary* for the expression of this supersensitivity.

Our results also show that dopamine supersensitive rats respond to D1 receptor agonists or antagonists as control rats do. Because our antipsychotic-treated rats showed a normal psychomotor response to a D1 partial agonist, this suggests that D1 receptor activation is not sufficient to trigger the expression of established dopamine supersensitivity. A D1 receptor antagonist also reduced the psychomotor response to d-amphetamine in haloperidol-pretreated rats, as it did in controls. Both findings are consistent with work showing that antipsychotic treatment does not produce consistent changes in striatal D1 receptor number (Fleminger et al., 1983; MacKenzie and Zigmond, 1985; Jiang et al., 1990; Marin and Chase, 1993). D-amphetamine also failed to increase ERK1/2 activity in the nucleus accumbens of dopamine-supersensitive rats. Because ERK activation by d-amphetamine requires D1 receptor activity (Valjent et al., 2005), this further supports the idea that the expression of dopamine supersensitivity following haloperidol treatment discontinuation does not involve potentiated D1-mediated neurotransmission in the nucleus accumbens.

Importantly, a D1 receptor antagonist normalized d-amphetamine-induced locomotion in haloperidol-pretreated rats, suggesting that D1-mediated neurotransmission mediates the expression of an exaggerated behavioural response to dopamine stimulation in these rats. This extends findings that chronic stimulation of D1 (but not D2) receptors reverses the expression of antipsychotic-evoked dopamine supersensitivity (Marin and Chase, 1993; Braun et al., 1997) [see also (Ramos et al., 2004; Shuto et al., 2006)]. As such, D1 receptors could be potential targets to temper the behavioural manifestations of supersensitivity. However, a caveat here is that D1 blockade also supressed basal locomotion in our rats, raising the possibility of non-d-amphetamine-specific motor effects. Using a lower dose of the D1 receptor antagonist SCH39166 or comparing its effects with other D1 receptor antagonists could resolve this issue. Another potential caveat here is that SKF38393 is a partial D1 agonist (Setler et al., 1978), and this could have reduced the likelihood of observing an enhanced psychomotor response in antipsychotic-treated rats. However, this is unlikely because SKF38393 dose-dependently increased psychomotor activity across groups (Fig. 4D), indicating that under our test conditions, SKF38393 had psychomotor activating effects. Future work can extend this finding using full D1 receptor agonists such as SKF81297 (Neumeyer et al., 2003).

### 4.2. Effects of different pro-dopaminergic agents

Following antipsychotic treatment cessation, dopamine-supersensitive rats showed an augmented psychomotor response to the monoamine releaser, d-amphetamine, the monoamine receptor agonist apomorphine and the D2 receptor agonist quinpirole. However, haloperidol-treated rats showed normal responses to the dopamine reuptake blocker GBR12783 and the monoamine reuptake blocker cocaine. Relative to these reuptake blockers, d-amphetamine’s unique psychomotor effects in dopamine-supersensitive rats could involve more potent actions at the dopamine transporter (DAT). For example, d-amphetamine enhances dopamine neurotransmission by both blocking dopamine uptake and enhancing dopamine release (Rothman et al., 2001) [but cocaine might also do this (Venton et al., 2006)]. However, *in vivo* microdialysis measurements show that d-amphetamine-induced increases in extracellular dopamine are unchanged in haloperidol-treated, dopamine-supersensitive rats (Samaha et al., 2007). D-amphetamine could also produce a supersensitive response in antipsychotic-treated rats through DAT-independent effects. For instance, in the caudate-putamen, d-amphetamine (but not cocaine) depletes dopamine-containing vesicles and enhances tonic dopamine release (Covey et al., 2013). It remains to be determined how antipsychotic-evoked dopamine supersensitivity might influence these processes.

### 4.3. Central processes

Much to our surprise, manipulations that increase central dopamine neurotransmission did not trigger the expression of established dopamine supersensitivity in rats previously treated with haloperidol. Administering neurotensin or DAMGO into the VTA—treatments that increase dopamine impulse flow (Kalivas and Duffy, 1990; Laitinen et al., 1990)—produced a similar psychomotor activating effect in dopamine-supersensitive and control rats. We did not measure neurotensin- or DAMGO-induced increases in extracellular dopamine concentrations directly. However, the concentrations we used increase both *i*) extracellular dopamine in terminal regions including the nucleus accumbens (Kalivas and Duffy, 1990; Laitinen et al., 1990), and *ii*) locomotion [Figs. 6C-D; see also (Kalivas and Duffy, 1990)], a highly dopamine-dependent behaviour. This suggests that increasing mesolimbic dopamine is not sufficient to evoke a supersensitive response in antipsychotic-treated rats. This could involve upregulated D2 autoreceptors in the VTA of these rats. When VTA D2 autoreceptors are stimulated by somatodendritic dopamine release, this reduces the excitability of local dopamine neurons. If chronic antipsychotic exposure upregulates D2 receptors in the VTA [as it does in the striatum (Burt et al., 1977; Fleminger et al., 1983)], dopamine activation of upregulated D2 autoreceptors could produce enhanced autoregulatory inhibitory feedback on dopamine neurons, thereby attenuating further dopamine release from both somatodendritic and terminal sites in antipsychotic-exposed rats. This in turn could explain why we did not observe enhanced locomotor responses to intra-VTA DAMGO or neurotensin in these rats. Another possible explanation is that rats with a history of antipsychotic exposure have reduced dopamine availability, such that neurotensin- or DAMGO-induced dopamine release is attenuated in these rats. However, this seems unlikely, because previous reports showed that chronic haloperidol exposure does not change striatal dopamine availability [(Compton and Johnson, 1988; Ichikawa and Meltzer, 1992; Samaha et al., 2007); see also (Demjaha et al., 2012; Howes et al., 2012)]. Nonetheless, it will be important to confirm and extend the present work with circuit-selective techniques such as chemogenetics and optogenetics.

It is also possible that injecting neurotensin or DAMGO into the VTA is not sufficient to trigger a sensitised response in dopamine-supersensitive rats, because peripheral actions are required to trigger this supersensitivity. In accord, injecting d-amphetamine into the cerebral ventricles produced a similar psychomotor response in dopamine-supersensitive and control rats. The lack of group differences could involve limited diffusion of d-amphetamine to relevant neural sites of action following intracerebroventricular infusion. However, we do not believe this to be the case. First, in control rats, intracerebroventricular d-amphetamine evoked a psychomotor response comparable in magnitude to that evoked by subcutaneous d-amphetamine (Fig. 7C). Moreover, infusing d-amphetamine directly into striatal subregions also produces comparable levels of locomotor activity in dopamine-supersensitive and control rats (El Hage et al., 2015). Together, the findings suggest that d-amphetamine’s effects in the brain are not sufficient to evoke a supersensitive response in antipsychotic-treated rats. This contrasts with observations that antipsychotic-treated rats show a sensitized locomotor response to intra-striatal dopamine infusions (Halperin et al., 1983, 1989). However, the antipsychotic doses used in these previous studies correspond to doses much higher than those used in the present work, and in humans (Kapur et al., 2003). Using a clinically-representative antipsychotic treatment regimen (Farde et al., 1989; Kapur et al., 2000; Kapur et al., 2003; Mamo et al., 2008), the present results suggest that in dopamine-supersensitive rats, the central effects of d-amphetamine are sufficient to increase psychomotor activity, but are not sufficient to evoke a supersensitive psychomotor response. Thus, the behavioural expression of antipsychotic-evoked dopamine supersensitivity might require peripheral actions. This can be studied further using dopamine antagonists that do not cross the blood-brain-barrier [such as domperidone (Laduron and Leysen, 1979)]. If d-amphetamine’s peripheral effects are necessary to trigger the expression of antipsychotic-evoked dopamine supersensitivity, a potential mechanism could involve the adrenal hormone corticosterone. Corticosterone is required for the expression of behavioural supersensitivity to d-amphetamine in other contexts. For instance, in rats, removing the adrenal glands supresses the exaggerated psychomotor response to d-amphetamine evoked by stress exposure, and this is restored by corticosterone replacement therapy (Deroche et al., 1992; Deroche et al., 1993). Greater circulating levels of corticosterone also predicts a greater psychomotor response to d-amphetamine (Piazza et al., 1991; Cador et al., 1993). In this context, examining the effects of chronic antipsychotic treatment and dopamine supersensitivity on corticosterone levels and function could be informative.

## 5. Conclusions

Developing effective treatments to prevent the expression of antipsychotic-evoked dopamine supersensitivity depends on a better understanding of the underlying biological mechanisms. In this context, our findings both extend existing knowledge on the role of D2 receptors in the expression of dopamine supersensitivity following antipsychotic treatment cessation and suggest two new underlying mechanisms (see Fig. 8). First, D1-mediated neurotransmission represents a potential target to temper the behavioural effects of antipsychotic-evoked dopamine supersensitivity. Second, while central dopamine neurotransmission (especially via D2 receptors) mediates the expression of antipsychotic-evoked dopamine supersensitivity, increasing central dopamine neurotransmission is not sufficient to trigger this behavioural response, suggesting that peripheral mechanisms are involved.

## Supporting information

Supplemental results (Figs. S1-S7)

## AUTHOR CONTRIBUTIONS

A. Servonnet: Conceptualization, methodology, validation, formal analysis, investigation, data curation, writing the original draft, reviewing and editing the manuscript and visualisation. F. Allain: Investigation and reviewing the manuscript. A. Gravel-Chouinard: Formal analysis and investigation. G. Hernandez: Investigation and reviewing the manuscript. C. Bourdeau Caporuscio: Investigation. M. Legrix: Investigation. D. Lévesque: Reviewing the manuscript and supervision. P-P. Rompré: Conceptualization, methodology and reviewing the manuscript. A-N. Samaha: Conceptualization, methodology, resources, reviewing and editing the manuscript, supervision, project administration and funding acquisition.

## Abbreviations

VTA: ventral tegmental area.

