## Supplemental results (Figs. S1-S7) for "Dopaminergic mechanisms underlying the expression of antipsychotic-induced dopamine supersensitivity in rats"

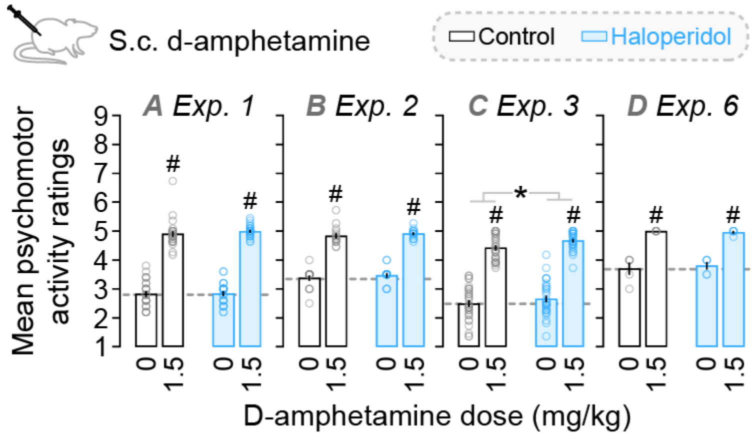


**Fig. S1. D-amphetamine effects on psychomotor activity ratings.** Across studies, d-amphetamine increased psychomotor activity ratings relative to vehicle (Injection effect; **A**, $F_{1,51} = 744.4$, $p < 0.0001$; **B**, $F_{1,30} = 502.4$, $p < 0.0001$; **C**, $F_{1,61} = 556.3$, $p < 0.0001$; **D**, $F_{1,8} = 99.86$, $p < 0.0001$). There were no group differences except in Exp. 3, where haloperidol rats had greater psychomotor activity ratings relative to controls (**C**; Group effect, $F_{1,61} = 5.03$, $p = 0.029$). Dotted lines indicate mean ratings of control rats receiving saline. *n*'s = 5-32/condition. $^{\#}p < 0.05$, relative to vehicle ('0 mg/kg') in the same group. $^{*}p < 0.05$. In (**C**), Group effect.

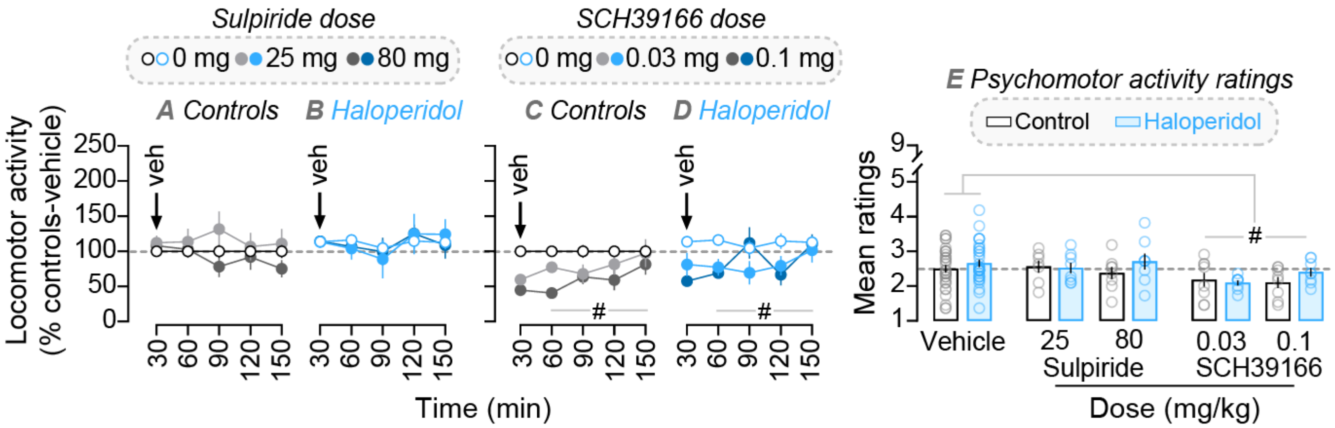


**Fig. S2. Effects of the D2 antagonist sulpiride and the D1 antagonist SCH39166 on vehicle-induced locomotion and psychomotor activity ratings.** In both haloperidol-treated and control groups, sulpiride had no influence on vehicle-induced locomotion (**A**-**B**) or on ratings (**E**, vehicle versus sulpiride) (all *P*'s > 0.05). SCH39166 reduced vehicle-induced locomotion and ratings similarly in haloperidol rats and controls (minutes 60-150; **C**-**D**; Injection × Time interaction, $F_{6,267} = 2.35$, $p = 0.03$; Injection effect, $F_{2,89} = 7.74$, $p = 0.001$; **E**; vehicle versus SCH39166; Injection effect, $F_{2,89} = 5.49$, $p = 0.006$). *n*'s = 7-32/condition. Dotted lines indicate response of control rats receiving vehicle. $^{\#}p < 0.05$. In (**C**-**D**), Injection × Time interaction and Injection effects. In (**E**), Injection effect.

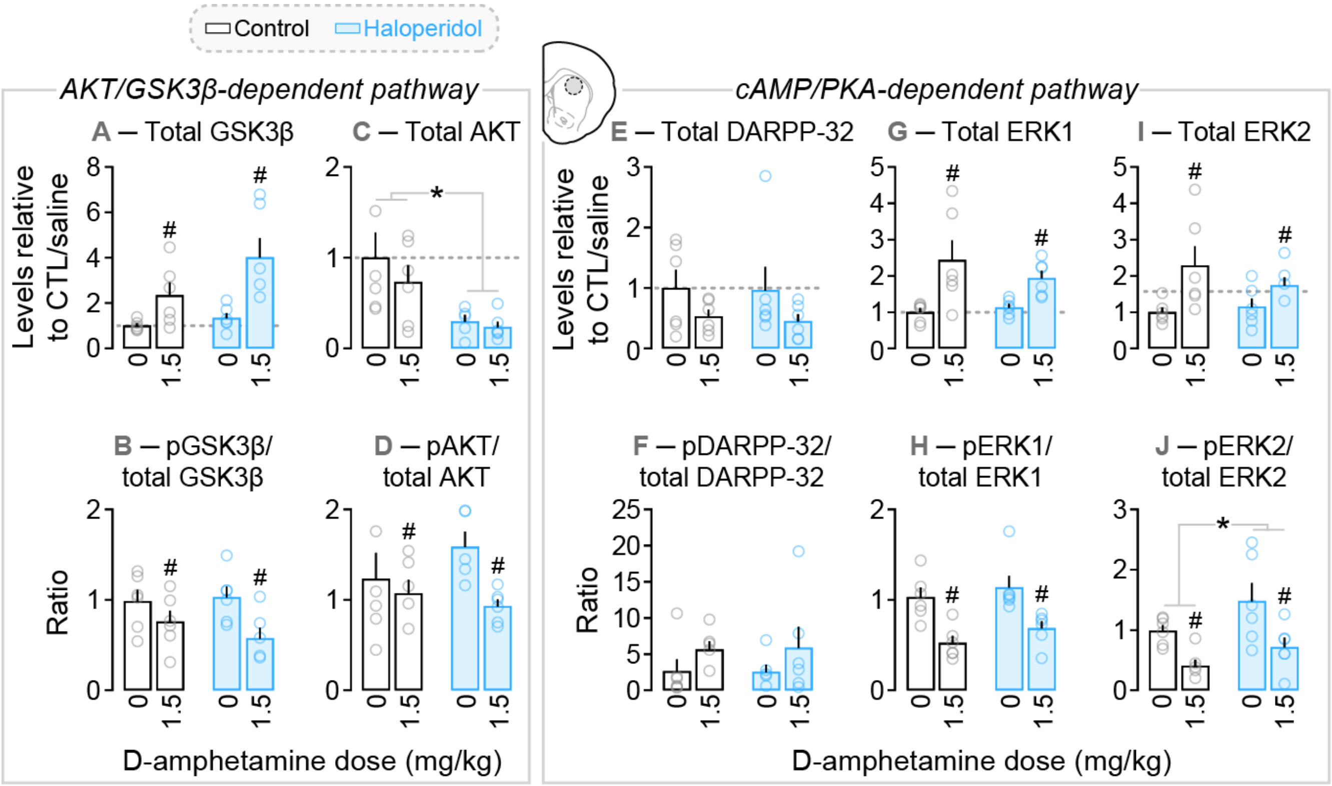


**Fig. S3. cAMP/PKA- and AKT/GSK3β-dependent signalling in the dorsal caudate-putamen of haloperidol-treated rats and control rats.** (**A**) Across groups, d-amphetamine increased GSK3β levels (Injection effect, $F_{1,20}$ = 15.19, $p$ = 0.0009), with no group differences. (**B**) D-amphetamine decreased pGSK3β/total GSK3β ratios similarly across groups (Injection effect, $F_{1,20}$ = 8.32, $p$ = 0.009). (**C**) Chronic haloperidol treatment decreased AKT levels, and this effect was similar after d-amphetamine or vehicle injection (Group effect, $F_{1,19}$ = 11.11, $p$ = 0.004). (**D**) Across groups, d-amphetamine decreased pAKT/total AKT ratios, with no group differences (Injection effect, $F_{1,19}$ = 4.65, $p$ = 0.04). There was no significant effect of haloperidol treatment or of d-amphetamine injection on (**E**) DARPP-32 or (**F**) pDARPP-32/total DARPP-32 ratios (all *P*'s > 0.05). Across groups, d-amphetamine enhanced (**G**) ERK1 and (**I**) ERK2 levels, and decreased (**H**) pERK1/total ERK1 and (**J**) pERK2/total ERK2 ratios (Injection effect; ERK1, $F_{1,20}$ = 14.65, $p$ = 0.0011; ERK2, $F_{1,20}$ = 9.05, $p$ = 0.007; pERK1/total ERK1 ratio, $F_{1,20}$ = 24.84, $p < 0.0001$; pERK2/total ERK2 ratio, $F_{1,20}$ = 13.87, $p$ = 0.0013). There were no group differences in these effects. (**J**) Prior haloperidol treatment increased pERK2/total ERK2 ratios, and d-amphetamine injection did not significantly change this effect (Group effect, $F_{1,20}$ = 4.94, $p$ = 0.038). *n*'s = 5-6/condition. In (**A-C-E-G-I**), dotted lines indicate protein levels in control rats injected with saline. $^{\#}p < 0.05$, relative to vehicle in the same group. $^{*}p < 0.05$. In (**C**, **J**), Group effect.

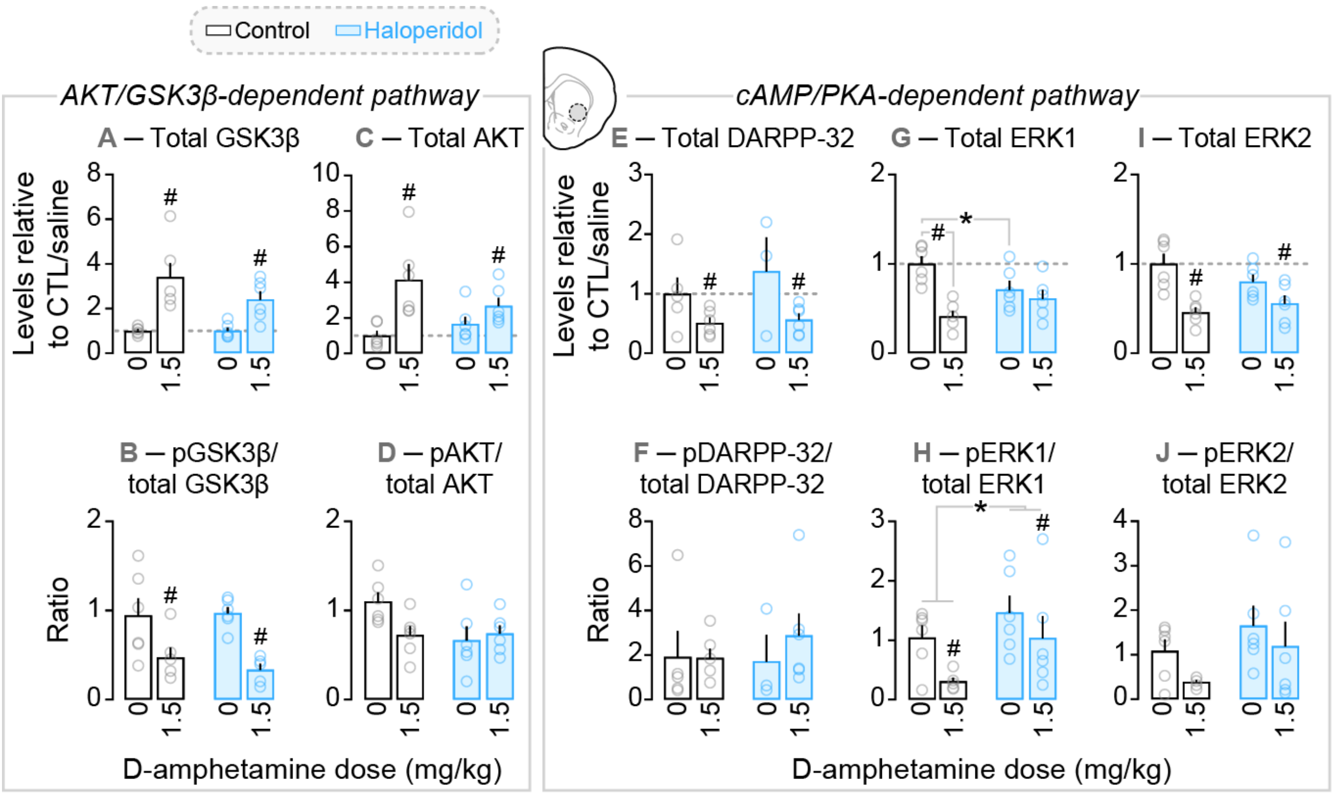


**Fig. S4. cAMP/PKA- and AKT/GSK3β-dependent signalling in the ventrolateral caudate-putamen of haloperidol-treated rats and control rats.** Across groups, d-amphetamine increased (**A**) GSK3β and (**C**) AKT levels (Injection effect; GSK3β, $F_{1,20} = 27.07$, $p < 0.0001$; AKT, $F_{1,20} = 14.35$, $p = 0.0012$), with no group differences. (**B**) D-amphetamine decreased pGSK3β/total GSK3β ratios similarly across groups (Injection effect, $F_{1,19} = 19.3$, $p = 0.0003$). (**D**) Neither haloperidol treatment nor d-amphetamine injection influenced pAKT/total AKT ratios ($p > 0.05$). Across groups, d-amphetamine decreased (**E**) DARPP-32 levels, (**I**) ERK2 levels and (**H**) pERK1/total ERK1 ratios (Injection effect; DARPP-32, $F_{1,16} = 8.11$, $p = 0.012$; ERK2, $F_{1,20} = 21.63$, $p = 0.0002$; pERK1/total ERK1 ratio, $F_{1,20} = 5.23$, $p = 0.033$), with no group differences. There was no effect of haloperidol treatment or of d-amphetamine injection on (**F**) pDARPP-32/total DARPP-32 ratios or (**J**) pERK2/total ERK2 ratios (all *P*'s > 0.05). (**G**) D-amphetamine decreased ERK1 levels in control rats relative to both vehicle in the same rats and vehicle in haloperidol rats (Group × Injection interaction, $F_{1,20} = 8.53$, $p = 0.0085$; Injection effect, $F_{1,20} = 17.04$, $p = 0.0005$; controls, saline > d-amph, $p = 0.0001$; saline, controls > haloperidol rats, $p = 0.05$). (**H**) Prior haloperidol treatment increased pERK1/total ERK1 ratios, and d-amphetamine injection did not influence this effect ($F_{1,20} = 5.07$, $p = 0.039$). *n*'s = 3-6/condition. In (**A-C-E-G-I**), dotted lines indicate protein levels in control rats injected with saline. $^{\#}p < 0.05$. In (**A**, **B**, **C**, **E**, **H**, **I**), relative to vehicle in the same group. $^{*}p < 0.05$. In (**H**), Group effect.

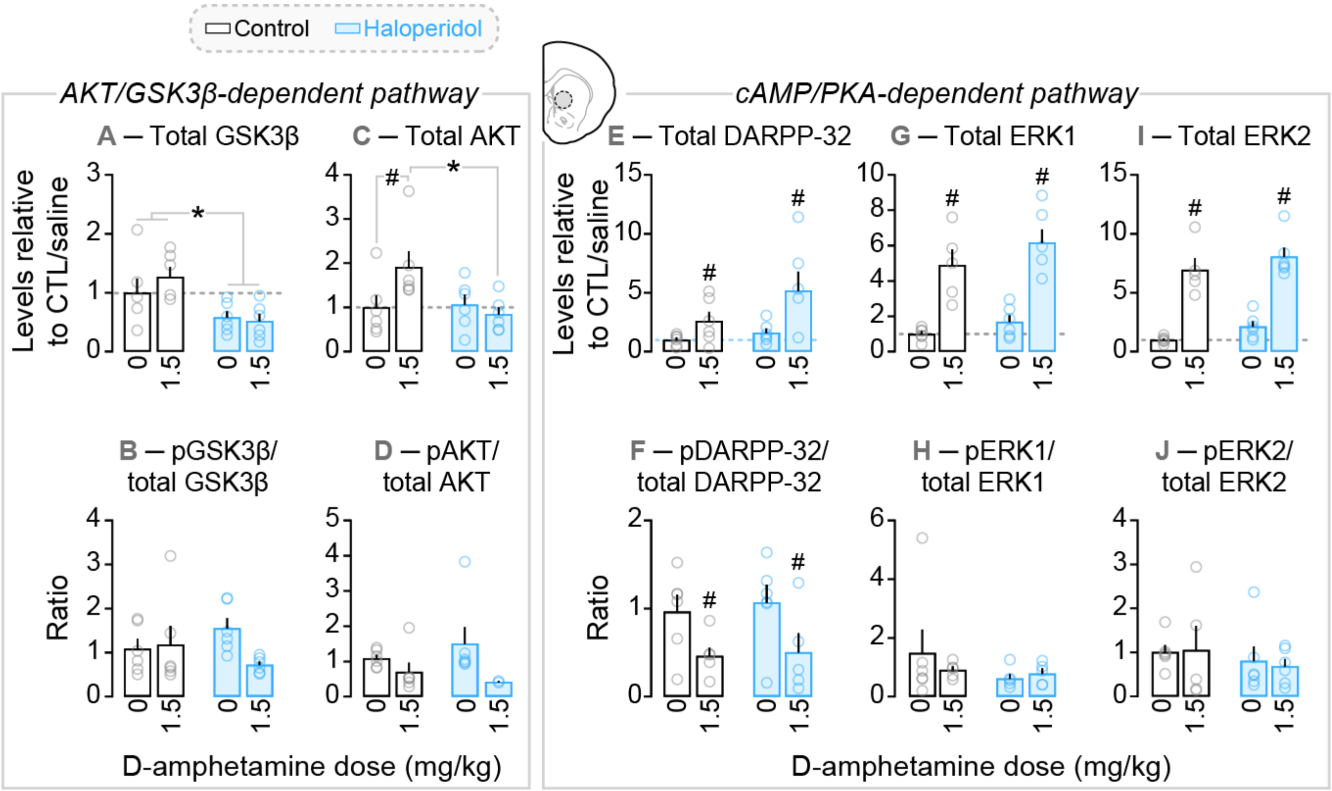


**Fig. S5. cAMP/PKA- and AKT/GSK3β-dependent signalling in the centromedial caudate-putamen of haloperidol-treated rats and control rats.** (**A**) Prior haloperidol decreased GSK3β levels, and this was not influenced by d-amphetamine injection (Group effect, $F_{1,20} = 12.49$, $p = 0.002$). There was no effect of haloperidol treatment or of d-amphetamine injection on (**B**) pGSK3β/total GSK3β and (**D**) pAKT/total AKT ratios (all $P$'s > 0.05). (**C**) D-amphetamine increased AKT levels in control rats only (Group × Injection interaction, $F_{1,20} = 4.75$, $p = 0.041$; controls, d-amph > saline, $p = 0.043$; d-amph, controls > haloperidol rats, $p = 0.017$). Across groups, d-amphetamine increased (**E**) DARPP-32, (**G**) ERK1 and (**I**) ERK2 levels (Injection effect; DARPP-32, $F_{1,20} = 8.03$, $p = 0.01$; ERK1, $F_{1,19} = 53.32$, $p < 0.0001$; ERK2, $F_{1,19} = 95.27$, $p < 0.0001$), with no group differences. (**F**) D-amphetamine decreased pDARPP-32/total DARPP-32 ratios similarly across groups (Injection effect, $F_{1,19} = 8.86$, $p = 0.008$). Neither prior haloperidol treatment nor d-amphetamine injection influenced (**H**) pERK1/total ERK1 ratios or (**J**) pERK2/total ERK2 ratios (all $P$'s > 0.05). $n$'s = 2-6/condition. In (**A**-**C**-**E**-**G**-**I**), dotted lines indicate protein levels in control rats injected with saline. $^{\#}p < 0.05$. In (**E**, **F**, **G**, **I**), relative to vehicle in the same group. $^{*}p < 0.05$. In (**A**), Group effect.

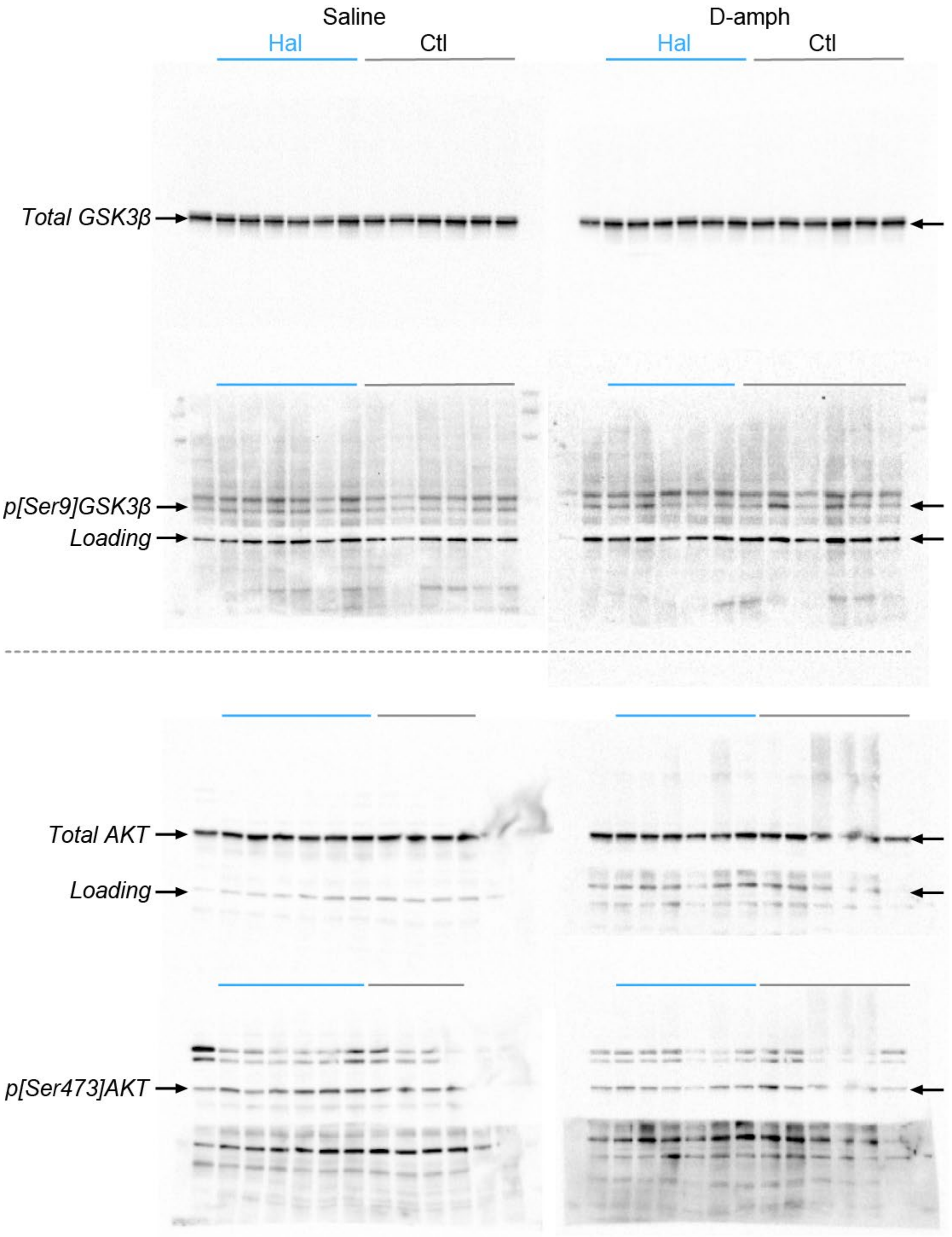


**Fig. S6. Western blots for GSK3β, p[Ser9]GSK3β, AKT and p[Ser473]AKT in nucleus accumbens tissue.** Uncropped pictures from Fig. 6. Note that loading control for AKT/p[Ser473]AKT was quantified on pictures with less light exposition.

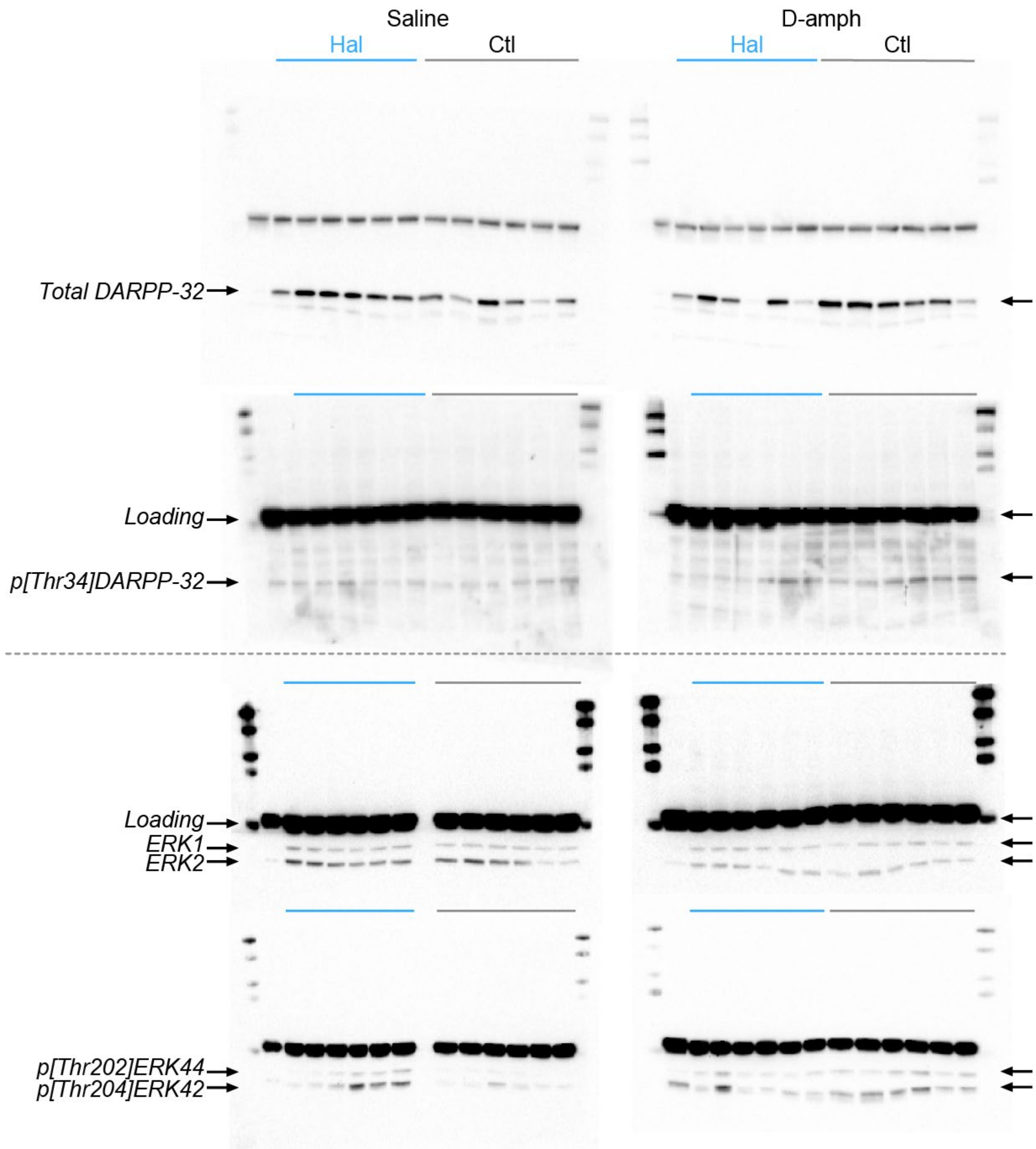


**Fig. S7. Western blots for DARPP-32, p[Thr34]DARPP-32, ERK1, ERK2, p[Thr202]ERK44 and p[Thr204]ERK42 in nucleus accumbens tissue.** Uncropped pictures from Fig. 6. Note that loading controls were quantified on pictures with more light exposition.
